# Insulin mRNA is stored in RNA granules in resting beta cells

**DOI:** 10.1101/2021.05.07.443159

**Authors:** Jovana Vasiljević, Djordje Vasiljević, Katharina Ganß, Anke Sönmez, Carolin Wegbrod, Esteban Quezada, Carla Münster, Eyke Schöniger, Daniela Friedland, Nicole Kipke, Marius Distler, Matthias Selbach, Michele Solimena

**Affiliations:** Molecular Diabetology, University Hospital and Faculty of Medicine Carl Gustav Carus, TU Dresden, Dresden, Germany; German Center for Diabetes Research (DZD e.V.), 85674 Neuherberg, Germany; Paul Langerhans Institute Dresden (PLID) of the Helmholtz Center Munich at the University Hospital Carl Gustav Carus and Faculty of Medicine of the TU Dresden, Dresden, Germany; Max Delbrück Center for Molecular Medicine (MDC), Berlin, Germany; Department of Visceral, Thoracic and Vascular Surgery, University Hospital and Faculty of Medicine Carl Gustav Carus, TU Dresden, Dresden, Germany; Charité – Universitätsmedizin Berlin, Berlin, Germany

**Keywords:** Beta cell, post-transcriptional regulation, RNA-binding proteins, stress granule, glucose-stimulated insulin biosynthesis

## Abstract

The glucose-stimulated biosynthesis of insulin in pancreatic islet beta cells is post-transcriptionally regulated. Several RNA-binding proteins (RBPs) that regulate *Insulin* mRNA stability and translation also bind mRNAs coding for other insulin secretory granule (ISG) proteins. However, an overview of these interactions and their glucose-induced remodelling is still missing. Here we identify two distinct sets of RBPs that were preferentially pulled down with the 5’-UTRs of mouse *Ins1*, *Ins2*, spliced *Ins2*, *Ica512*/*Ptprn* and *Pc2*/*Pcsk2* mRNAs from extracts of either resting or stimulated mouse insulinoma MIN6 cells. Among RBPs binding to all tested transcripts in resting conditions was hnRNP A2/B1. *Hnrnpa2b1* KO MIN6 cells contained lower levels of *Ins1* mRNA, proinsulin and insulin, and had reduced insulin secretion. In resting cells, both hnRNP A2/B1 and *Insulin* mRNAs localized to stress granules, which dissolved upon glucose stimulation. *Insulin* mRNA-positive RNA granules were also found in human pancreatic beta cells *in situ*. Our results suggest that resting beta cells store mRNAs for insulin secretory granule proteins in stress granules through specific RNA protein interactions. Glucose stimulation remodels these interactions, releasing the transcripts, and another set of RBPs coordinates their translation.

## Introduction

Insulin is the main hormone for control of blood glucose homeostasis. It is produced in pancreatic beta cells and packaged into secretory granules that undergo glucose-stimulated secretion. After a meal, the increase in blood glucose concentration prompts beta cells to secrete insulin into the bloodstream. Insulin stimulates glucose uptake into muscle and fat cells, while it inhibits gluconeogenesis and glucose release of hepatocytes, hence lowering blood glucose concentration to fasting levels. The beta cells, therefore, have one main function: to produce and secrete enough insulin in order to regulate blood glucose surges.

Surprisingly, beta cells secrete <5% of their insulin stores upon glucose stimulation^1^. On the other hand, newly-synthesized insulin is preferentially secreted^2^. Glucose stimulates *de novo* insulin biosynthesis and secretory granule biogenesis at a lower threshold than that for glucose-stimulated insulin secretion^3^, which maintains optimal stores of “new” secretory granules^4^. Impaired glucose responsiveness of beta cells, such as reduced ability to upregulate secretory granule production in response to insulin resistance and hyperglycemia may lead to reduced insulin secretion and type 2 diabetes (T2D). On the other hand, a moderately leaky biosynthesis and release of insulin in fasting conditions may also drive insulin resistance, and thus the vicious circle leading to T2D^5^.

*Insulin* mRNA accounts for up to 30% of beta cell transcriptome^6^, and each beta cell produces on average >3 x 10^3^ preproinsulin peptides every second^7^. This outstanding translation rate comes with several challenges. First, preproinsulin translation needs to increase quickly upon glucose stimulation in order for the younger secretory granules to be rapidly replenished. Accordingly, insulin biosynthesis is specifically regulated: while stimulated beta cells increase total protein biosynthesis ∼2-fold, they enhance that of preproinsulin up to 20-fold^8, 9^. Also, the newly-synthesized insulin must be packaged into newly-assembled secretory granules, which contain numerous other proteins required for proinsulin conversion and its exocytosis^10–12^. Among these are proprotein convertases PC1/3 and PC2, islet cell autoantigen ICA512 (also known as IA-2 and PTPRN), members of the granin family and SNARE proteins. In order for beta cells to operate properly, they must quickly and specifically coordinate the translation of many secretory granule proteins.

Glucose-stimulated beta cells increase insulin biosynthesis, even while the mRNA levels remain unchanged for up to at least 2 hours after stimulation^9, 13, 14^, implying the involvement of post-transcriptional mechanisms. Other proteins of insulin secretory granules are similarly post-transcriptionally regulated, such as PC1/3, PC2, ICA512 and granins^8, 15, 16^.

Post-transcriptional regulation relies on the binding of RNA-binding proteins (RBPs) to regulatory sequences in mRNAs, together forming ribonucleoprotein (RNP) complexes^17, 18^. It is the unique RNP code that confers gene-specific post-transcriptional regulation and defines whether a mRNA will be transported, translated, repressed and stored, or degraded^19^. These processes take place in membraneless organelles called RNA granules. Each different type of RNA granules serves a specific function: repressed mRNAs are stored in stress granules upon cellular stress, whereas mRNA degradation takes place in P bodies^20–22^. Remodelling of RNP complexes alters the fate and localization of mRNAs and thereby modifies the gene expression profile^18, 19^. Such post-transcriptional regulation has many advantages. Cells can quickly activate protein biosynthesis without depending on *de novo* transcription. Also, different mRNAs can be diversely regulated by different RBPs, whilst functionally related mRNAs can be co-regulated through sequences recognized by the same set of RBPs.

Pancreatic islet beta cells exploit these mechanisms to fine-tune their control of glucose homeostasis. It has been proposed that glucose activates the presynthesized, inactive *Insulin* mRNA^23^, presumably by remodelling the RBPs bound to its regulatory sequences. Within the 5’-UTR of *Insulin* mRNA, a 29 nucleotide (nt) stem loop^24^ and the preproinsulin glucose element (ppIGE)^25^ are required for its glucose-induced increase in translation. Enhanced stability of *Insulin* mRNAs is conveyed by a polypyrimidine motif^6, 16^ and a *UUGAA* motif at its 3’-UTR^9^. These sequences are conserved in rat, mouse and human *Insulin* transcripts^6, 26, 27^, and some of them are conserved in the UTRs of mouse *Pc1/3, Pc2*, *Ica512, pro-islet amyloid polypeptide* and *chromogranin A*^28^ mRNAs.

Some *Insulin* mRNA RBPs have been previously identified. The best characterized is polypyrimidine-tract binding protein 1 (PTBP1), which coordinates the expression of multiple secretory granule cargoes^27^. Glucose-stimulated binding of PTBP1 to a polypyrimidine sequence in the 3’-UTR of rat *Insulin1* mRNA increases the stability of the latter ^6^. Our previous results indicate that Ptbp1 similarly binds and stabilizes the 3’-UTRs of the rat *Insulin2*, *Pc2* and *Ica512* mRNAs upon glucose stimulation^16, 29^. Additionally, PTBP1 binding to the 5’-UTRs of these mRNAs stimulates their cap-independent translation, thereby selectively increasing the expression of secretory granule proteins^30^.

RBPs are among the most rapidly regulated classes of proteins in glucose and IBMX treated INS-1 cells^31^, suggesting their critical role for rapid responses of beta cells to glucose. Furthermore, nuclear retention of PTBP1^32^ and its suppression following exposure of human islets to prolonged hyperglycemia^33^ reduces glucose-stimulated insulin biosynthesis and secretion, respectively. Common polymorphisms in *PTBP1* are linked to impaired glucose stimulated insulin secretion in humans^34^.

While it is accepted that glucose remodels RBPs bound to mRNAs for insulin and other secretory granule proteins, we lack a global understanding of the RNP code in different conditions. Additional RBPs that post-transcriptionally regulate insulin expression include HuD^35, 36^, TIAR^37^, DDX1^38^, however, their involvement in co-regulating the expression of several insulin secretory granule proteins is unknown. Furthermore, the storage of the presynthesized *Insulin* mRNA in the cytoplasm was vaguely proposed^23^, but the details are not known.

Here, we used *in vitro* RNA pull-downs coupled with mass spectrometry (MS) to identify factors binding to the 5’-UTRs of mRNAs for several insulin secretory granule proteins. We show that these mRNAs share several common interacting RBPs, which differ in resting and glucose stimulated conditions. We confirmed that one of these proteins, hnRNP A2/B1, regulates insulin production. We also demonstrate that the *Insulin* mRNA is located in stress granules in resting cells, which dissolve upon glucose stimulation. We propose that in resting beta cells, stress granules store and repress transcripts for insulin secretory granule proteins by binding to a set of co-regulating RBPs. Glucose entry into beta cells fosters the remodelling of RNP complexes, thereby causing the dissolution of stress granules and enabling a co-regulated burst in mRNA translation and insulin secretory granule biogenesis.

## Results

### The 5’UTRs of secretory granule proteins share common protein binders in resting and glucose-stimulated conditions

Several proteomic approaches enable identification of proteins interacting with individual mRNA sequences^39^. In order to determine if glucose remodels RNPs regulating the translation of insulin secretory granule proteins, we performed *in vitro* RNA pull-downs using cytoplasmic extracts of resting and stimulated mouse insulinoma MIN6 cells, and identified the bound proteins using MS (fig. 1a). We chose the 5’-UTRs of mouse *Ins1*, *Ins2*, *Pc2* and *Ica512* mRNAs as target transcripts, since these are known to be post-transcriptionally coordinated by PTBP1^30^ and hnRNP K (our unpublished data). In mouse beta cells the majority of insulin results from the translation of *Ins2* mRNA^40^. At variance with mouse islets, in our MIN6 cells *Ins2* mRNA is twice as abundant as *Ins1* mRNA. Because this shorter transcript variant is spliced within its 5’-UTR, we also included it in our assays.

**Figure 1.**
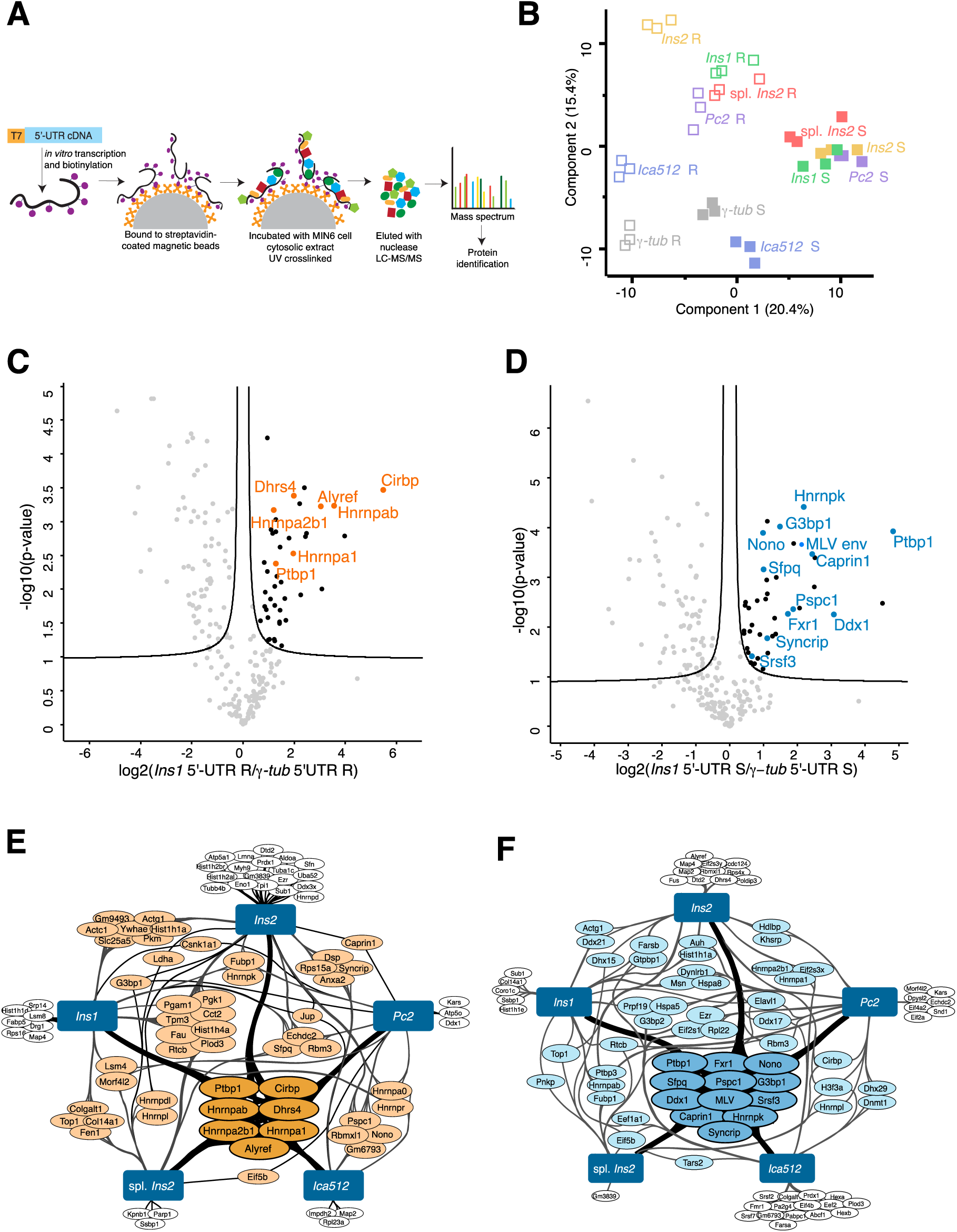
A specific set of RBPs bind to 5’-UTRs of mRNAs encoding secretory granule proteins in resting and in stimulated MIN6 cells. **A:** cartoon of *in vitro* RNA pull-down. **B:** PCA plot shows separate clustering of secretory granule protein mRNPs from cytosolic extracts of resting (R, open squares) and stimulated (S, filled squares) MIN6 cells. The *γ-tub* 5’-UTR was used as a control. **C and D:** volcano plots showing significantly enriched proteins binding to the *Ins1* 5’-UTR compared to the *γ-tub* 5’-UTR in resting (orange, **C**) and stimulated (blue, **D**) MIN6 cells. **E and F:** interaction plots of all significantly enriched RBPs that bind to the secretory granule protein mRNA 5’-UTRs in resting (orange, **E**) and stimulated (blue, **F**) MIN6 cells. Proteins that were shared among all transcripts are labeled with a darker color. Ins – insulin; Pc2 – prohormone convertase 2; Ica512 – islet cell autoantigen 512; γ-tub – γ-tubulin; R – resting, 2 h, 0 mM glucose; S – stimulated, 25 mM glucose, 2h.

To identify RBPs that specifically bind to the target 5’-UTRs, we required a control to account for unspecific binding. Since the expression of our chosen insulin secretory granule proteins is glucose regulated, we assumed that their transcripts bind RBPs that do not equally bind to transcripts for housekeeping proteins or to common RNA features, such as the 5’ cap. The mouse *γ-tubulin* mRNA 5’-UTR did not bind PTBP1 and hnRNP K, our positive RBP controls (suppl. fig. 1a). Therefore, the 5’-UTR of the mouse *γ-tubulin* mRNA was used as a control. The identified proteins and log2 transformed LFQ values are shown in **suppl. table 1**, and data are available via ProteomeXchange with identifier PXD026956.

Fig. 1b shows a clear segregation of our samples according to glucose stimulation with replicate samples clustering together, indicating high reproducibility. Moreover, based on their interacting proteins, RNP complexes for secretory granule proteins cluster separately from those for *γ-tubulin* in both resting and stimulated conditions. Replica experiments displayed a similar PCA profile (suppl. fig. 1b). This was true for all tested 5’-UTRs except for that of *ICA512* mRNA. These results corroborate the hypothesis that a specific RNP code enables the selective glucose-dependent regulation of mRNAs for insulin secretory granule proteins.

Since the RNPs of transcripts coding for secretory granule proteins formed separate clusters in resting and glucose-stimulated conditions, we assumed that the same RBPs bind to these transcripts. With permutation-based t-tests, we compared the enrichment of the proteins interacting with each target mRNA 5’-UTR to that of *γ-tubulin* in the same condition (fig. 1c–d and suppl. fig. 1c-d). Significance was determined as described in ^41^, with FDR = 0.05 and slope s0 = 0.1 (**suppl. table 2**). In resting conditions, we recovered seven RBPs selectively enriched with all target 5’-UTRs (labelled in orange, fig. 1c and 1e, and suppl. fig. 1c). Similarly, in stimulated conditions, there were twelve common significantly enriched RBPs (labelled in blue, fig. 1d and 1f, and suppl. fig. 1d). PTBP1 was specific for all of the target RNPs in both resting and glucose-stimulated conditions, whereas hnRNP K was a shared RBP enriched upon stimulation. DDX1, which was shown to interact with the *Insulin* mRNA in rat insulinoma INS-1 cells^38^, was specific for the target sequences upon stimulation. Hence, we identified a common set of RBPs associated with secretory granule protein transcripts in resting conditions that are remodelled upon glucose stimulation. To our knowledge, apart from PTBP1, DDX1 and hnRNP K, the other common RBPs, such as G3BP1 and several members of the hnRNP family, have not previously been associated with secretory granule protein mRNAs.

### hnRNP A2/B1 binds to the 5’-UTRs of mRNAs for secretory granule proteins

hnRNP A2/B1 is one of the identified RBPs that displayed significantly enriched binding to secretory granule protein transcripts in resting conditions (fig. 1c and 1e, and suppl. fig. 1c). As a member of the hnRNP family, hnRNP A2/B1 functions in virtually all aspects of post-transcriptional regulation^42, 43^. Previously, hnRNP A2/B1 has been shown to reside in RNA granules containing repressed RNAs, such as transport granules and stress granules neurons and oligodendrocytes^44–47^. Since we hypothesized that resting beta cells store secretory granule protein transcripts in the cytoplasm, we further investigated the role of hnRNP A2/B1 in regulating the production of insulin and other secretory granule proteins.

We first validated the binding of hnRNP A2/B1 to mRNAs for secretory granule proteins by RNA-IP. The recovery of *Ins1*, *Ins2* and *Pc2* mRNAs upon hnRNP A2/B1 immunoprecipitation (IP) was enriched compared with the recovery of *γ-tubulin* mRNA (fig. 2a), while *Ica512* mRNA recovery was similar to that of *γ-tubulin* mRNA (fig. 2a), possibly reflecting their positions on the PCA plot (fig. 1b).

**Figure 2.**
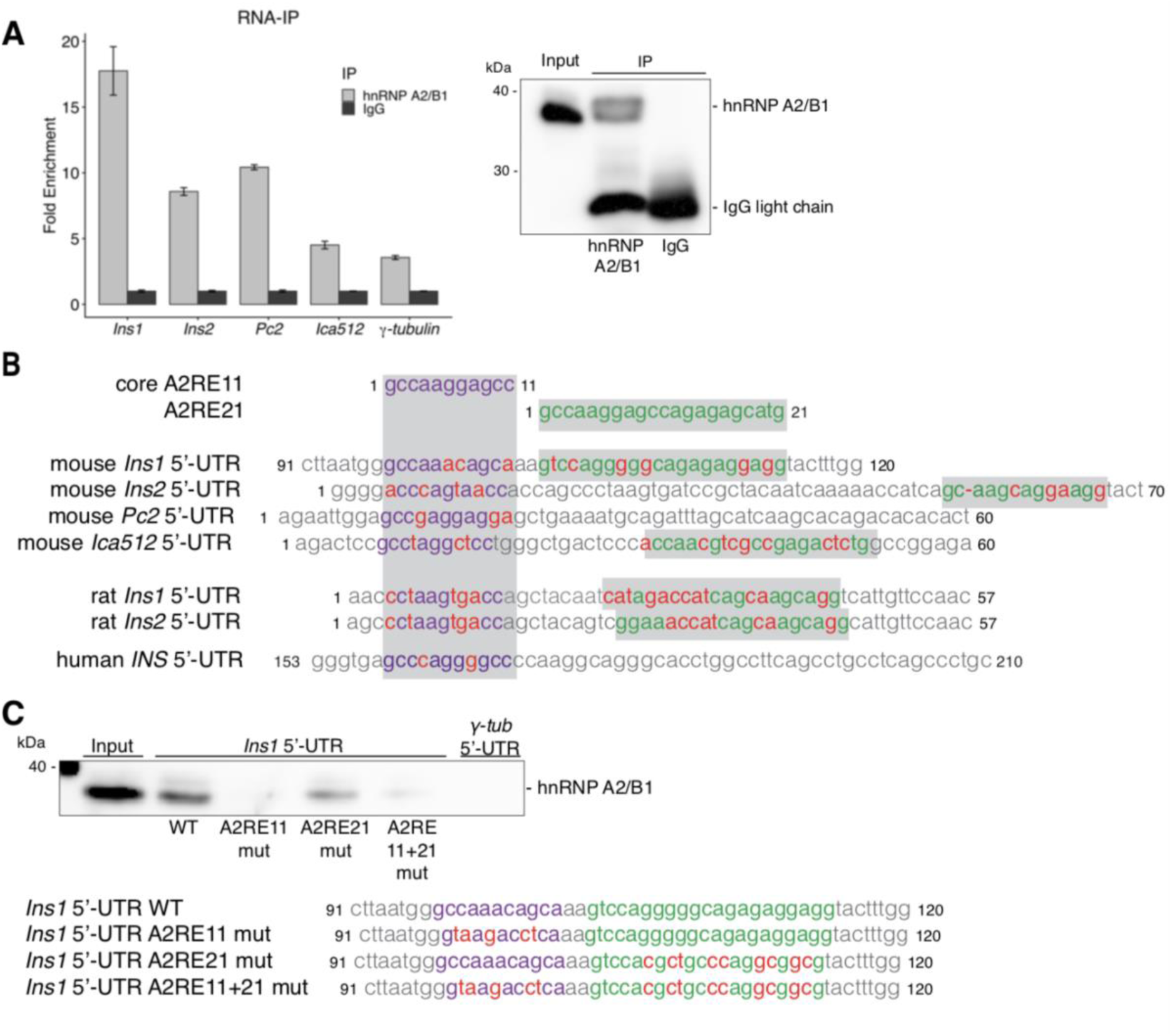
hnRNP A2/B1 binds to the 5’-UTRs of secretory granule protein transcripts. **A:** co-purification of secretory granule protein transcripts by hnRNP A2/B1 immunoprecipitation (IP) and quantified by qPCR. Fold enrichment was calculated compared to a nonspecific IgG IP (n=3, mean±SD). Panel on the right shows a representative image demonstrating antibody specificity. **B:** sequence and location of putative A2REs in the 5’-UTRs of mouse secretory granule protein transcripts, and human and rat insulin transcripts. Core A2RE11 labeled in purple and longer A2RE21 labeled in green, nucleotides differing to the consensus sequence labeled in red. **C:** reduced binding of hnRNP A2/B1 to the *Ins1* 5’-UTR with A2RE mutations. Mutated nucleotides shown below in red. Ins – insulin; Pc – prohormone convertase; Ica512 – islet cell autoantigen 512; γ-tub – γ-tubulin; A2RE – A2 response element.

The binding sequence of hnRNP A2/B1 consists of a 21 nucleotide (nt) long A2 response element (A2RE21), with a shorter A2RE11 defined as the core binding motif^44, 48^. We identified sequences homologous to the A2REs within the 5’-UTRs of several secretory granule protein transcripts (fig. 2b). These sequences are conserved in mouse, rat and human insulin transcripts. In most of these transcripts, the A2RE11 was positioned first, followed by the longer A2RE21. We tested if hnRNP A2/B1 binds to these sequences by RNA pull-down with the *Ins1* mRNA 5’-UTR harboring mutated A2REs (fig. 2c). The mutation of the A2RE11 both individually and in combination with the A2RE21 abolished hnRNP A2/B1 binding to the *Ins1* mRNA 5’-UTR, while its binding upon A2RE21 mutation was reduced. These data indicate that hnRNP A2/B1 binds to *Ins1* mRNAs through the A2REs in its 5’-UTR. Presumably, the putative A2REs in the other secretory granule protein transcripts also specifically bind hnRNP A2/B1.

### hnRNP A2/B1 regulates the expression of secretory granule proteins

Since we determined that hnRNP A2/B1 binds to the 5’-UTRs of secretory granule protein mRNAs through A2REs, we investigated whether it affects their expression. We generated an *Hnrnpa2b1* KO MIN6 cell clone using CRISPR/Cas9 editing with a guide positioning the cleavage near the start codon (suppl. fig. 2a, boxed region). The editing abolished *Hnrnpa2b1* expression due to a large insertion that removed its start codon (suppl. fig. 2b and 2c). While *Ins1* mRNA levels were reduced in both resting and glucose-stimulated *Hnrnpa2b1* KO cells, *Ins2* mRNA levels were unchanged (fig. 3a). Glucose-stimulated *Hnrnpa2b1* KO cells contained less insulin compared to WT cells (fig. 3b), while proinsulin, as measured by ELISA, was reduced in both resting and stimulated *Hnrnpa2b1* KO cells (fig. 3c), indicating that hnRNP A2/B1 regulates insulin biosynthesis regardless of glucose concentrations. Moreover, ELISA confirmed the lower insulin content of *Hnrnpa2b1* KO cells, which secreted less insulin upon stimulation (fig. 3c). However, the stimulation index of WT and *Hnrnpa2b1* KO cells did not differ, indicating that their secretory capacity was not altered and that the reduced insulin release reflects lower insulin production in these cells.

**Figure 3.**
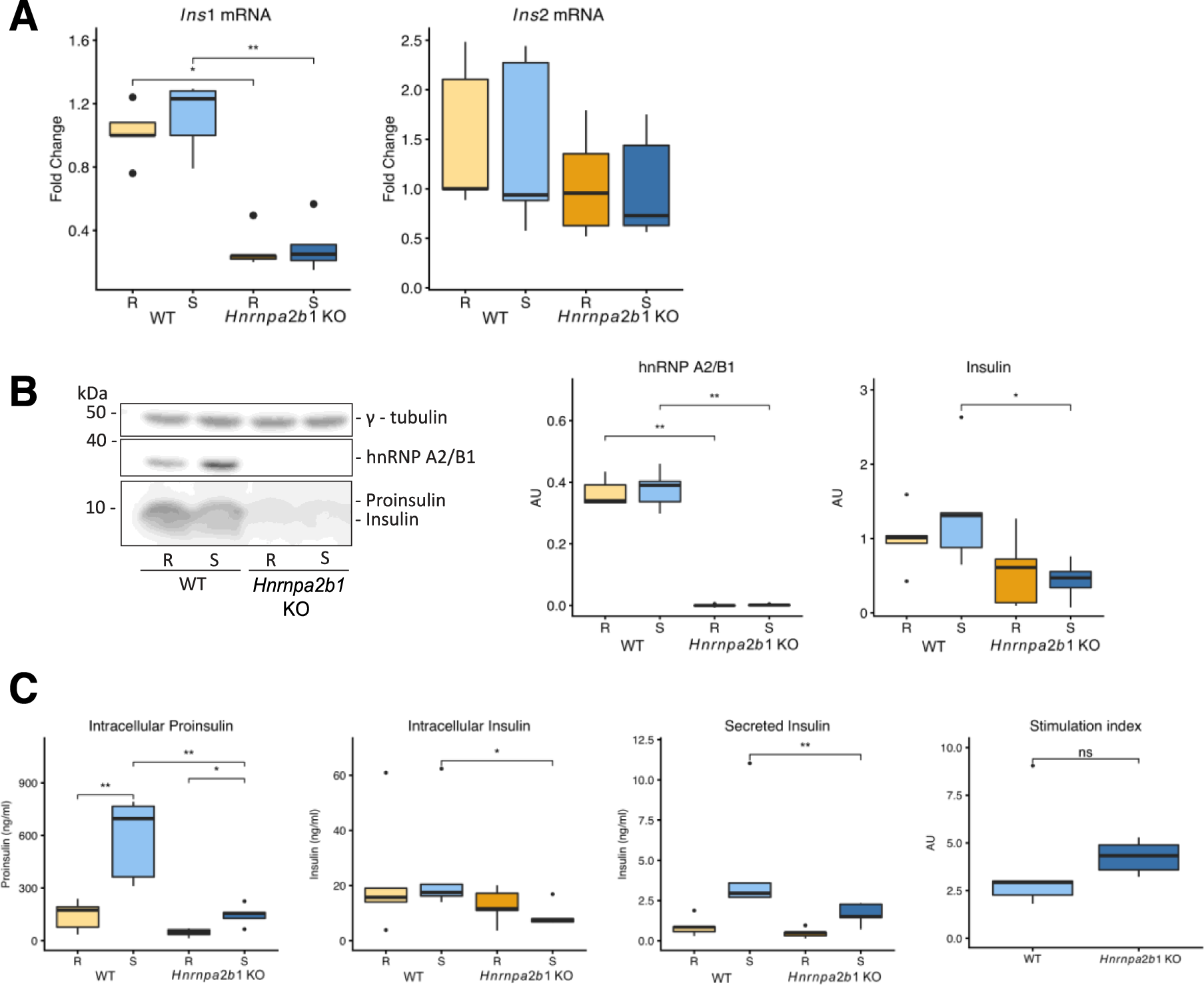
hnRNP A2/B1 KO reduces insulin expression in MIN6 cells. **A:** reduction of *Ins1* but not *Ins2* RNA in resting and stimulated *Hnrnpa2b1* KO MIN6 cells, normalized to *β-actin* mRNA. *Ins1* mRNA is reduced in *Hnrnpa2b1* KO cells. **B:** hnRNP A2/B1 and insulin protein levels are reduced in resting and stimulated *Hnrnpa2b1* KO MIN6 cells. Normalized to γ-tubulin. **C:** intracellular proinsulin and insulin, and secreted insulin are reduced in stimulated *Hnrnpa2b1* KO MIN6 cells compared to WT cells, as measured by ELISA and normalized to DNA concentration. Stimulation index, calculated as secreted insulin divided by total insulin and normalized to resting insulin levels, is unchanged. Ins – insulin; R – resting, 2 h, 0 mM glucose; S – stimulated, 25 mM glucose, 2h. Plots show Tukey-style boxplots of five independent replicates. Mann-Whitney test: * = p<0.05, ** = p<0.01.

Even though hnRNP A2/B1 binds to the 5’-UTRs of *Pc2* and *Ica512* mRNAs, the levels of these transcripts in *Hnrnpa2b1* KO and WT cells were similar (suppl. fig. 2d). Moreover, glucose stimulation similarly upregulated the levels of PC1/3, PC2 and ICA512 proforms in WT and *Hnrnpa2b1* KO cells (suppl. fig. 2e). Possibly, PC1/3, PC2, and ICA512 expression in the *Hnrnpa2b1* KO cells can be rescued by a paralogue protein with redundant functions. One possible candidate is hnRNP A1, which is also found in the secretory granule protein RNP complexes in resting conditions (fig. 1c and 1e, and suppl. fig. 1c) and is upregulated in *Hnrnpa2b1* KO cells (suppl. fig. 2e).

### hnRNP A2/B1 regulates the translation of secretory granule proteins in beta cells

We have shown that hnRNP A2/B1 regulates insulin production, but the mechanism of action is unclear. To further explore its role in insulin translation, we generated several translation reporters to test if the mutation of A2REs in the *Ins1* mRNA 5’-UTR affected the biosynthesis of a downstream luciferase reporter (fig. 4b). Compared with the WT sequence, we measured reduced luciferase activity in reporter constructs harboring mutated A2REs. The mutation of A2RE21 in the *Ins1* mRNA 5’-UTR resulted in the lowest luciferase signal, which was similar to the signal detected with the *γ-tubulin* mRNA 5’-UTR coupled luciferase construct. The upregulation in glucose-stimulated insulin biosynthesis is 10 times more than the upregulation of the total beta cell proteome^8^. Our data demonstrate that the binding of hnRNP A2/B1 contributes to this specific regulation. We suggest that hnRNP A2/B1 is an RBP that specifically regulates the expression of *Insulin* mRNA relative to other housekeeping genes.

**Figure 4.**
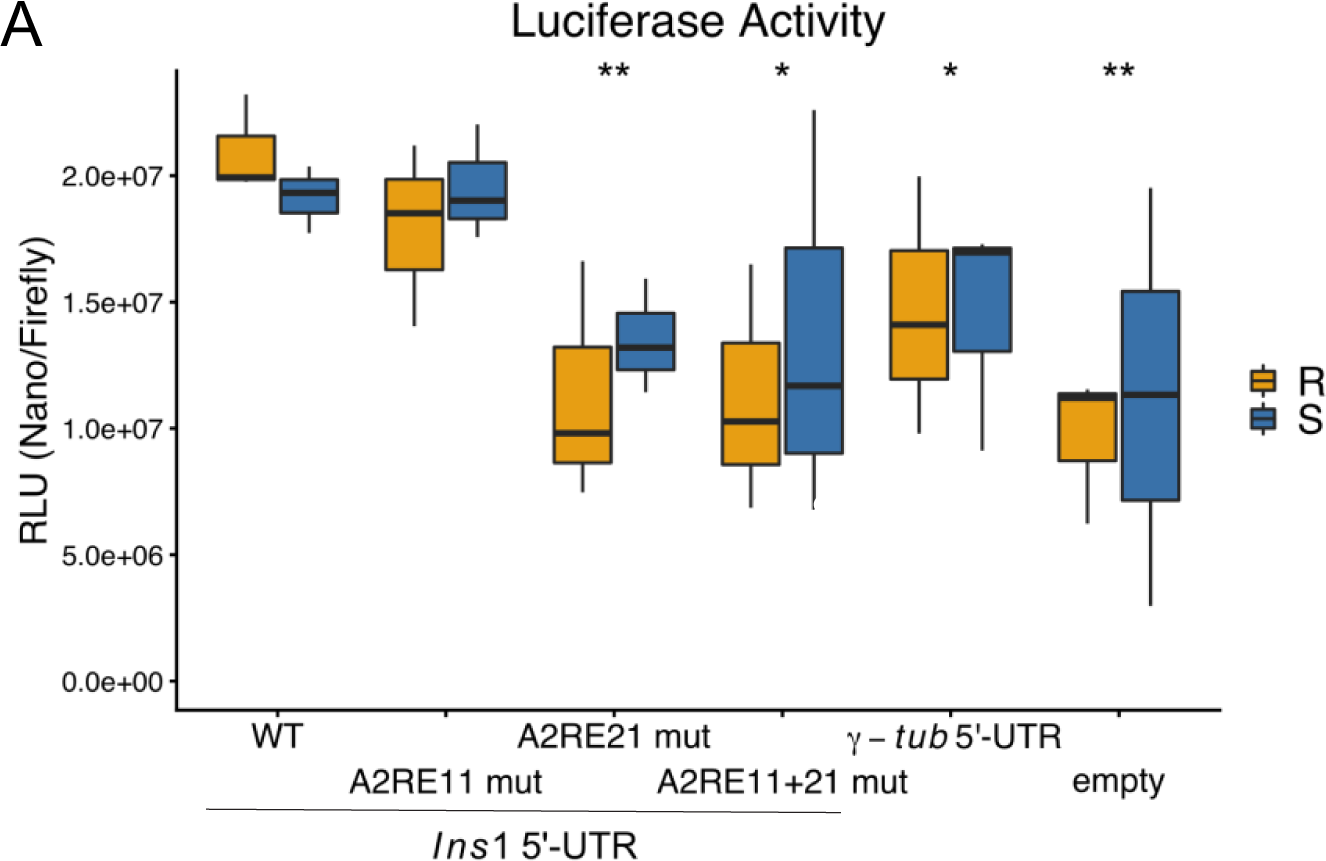
hnRNP A2/B1 controls insulin biosynthesis MIN6 cells. **A:** Nanoluciferase activity of constructs with mutated A2REs the *Ins1* mRNA 5’-UTR is reduced compared to WT levels, normalized to firefly, mutations as in fig. 2C. Plots show Tukey-style boxplots of three independent replicates. Mann-Whitney test: * = p<0.05, ** = p<0.01. Ins – insulin; γ-tub – γ-tubulin; A2RE – A2 response element.

### hnRNP A2/B1 and *Insulin1/2* mRNAs reside in stress granules in the cytoplasm of resting MIN6 cells

hnRNP A2/B1 is a component of RNA granules that contain repressed mRNAs^44–47^. For this reason, we investigated its localization in resting and glucose stimulated MIN6 cells. In resting cells hnRNP A2/B1 was localized to cytoplasmic structures which resemble RNA granules, which were absent in glucose stimulated cells (suppl. fig. 3a). RNAse treatment of the cells prior to fixation abolished the presence of these structures, confirming their identity as RNA granules (suppl. fig. 3b).

We further verified the identity of hnRNP A2/B1^+^ structures with different RNA granule markers. In resting cells, hnRNP A2/B1 co-localized with stress granule markers eIF3b and G3BP1 (suppl. fig. 3c and 3d), but not with the P-body marker Dcp1a (suppl. fig. 3e). The colocalization of G3PB1 and eIF3b was used as a positive control (suppl. fig. 3f), and colocalization was quantified using the Pearson’s correlation coefficient (suppl. fig. 3g). Both stress granule markers were highly colocalized with hnRNP A2/B1 in resting, but not in glucose stimulated cells. As stress granules are compartments where repressed mRNAs are stored during cell stress, we hypothesized that resting beta cells use hnRNP A2/B1 ^+^ RNA granules to store *Ins1/2* mRNA and protect it from degradation.

Since hnRNP A2/B1 binds to the 5’-UTRs of mRNAs coding for secretory granule proteins (fig. 1c, 1e, 2a and suppl. fig. 1c), we presumed that these transcripts colocalize with hnRNP A2/B1^+^ stress granules. In resting MIN6 cells, *Ins1/2* mRNA colocalized indeed with hnRNP A2/B1^+^ granules (suppl. fig. 4a). This colocalization was lost upon glucose stimulation, where the *Ins1/2* mRNA was dispersed throughout the cytoplasm (suppl. fig. 4a), presumably being translated at the ER. To verify the specificity of the probes, we confirmed that mouse glucagonoma α-TC cells were negative for *Ins1/2* mRN*A* and positive for the control *Gapdh* mRNA (suppl. fig. 4b). Additionally, we verified that *Gapdh* mRNA does not localize to stress granules in resting MIN6 cells (suppl. fig. 4c). Taken together, these data suggest that resting MIN6 cells store *Ins1/2* mRNAs in stress granules. Glucose stimulation remodels RBPs bound to these mRNAs and dissolves stress granules, releasing the mRNAs for translation.

A caveat of the MIN6 cells used in all previous experiments is their threshold of 1-2 mM glucose for stimulated insulin biosynthesis and secretion, which is much lower than the 5-5.5 mM glucose threshold of primary mouse beta cells. Accordingly, 2.8 mM glucose already activates the biosynthesis of secretory granule proforms in these MIN6 cells (suppl. fig. 4d), which therefore only rest in media with 0 mM glucose. However, as glucose absence is equivalent to starvation, the stress granules detected in resting MIN6 cells may reflect a stress response rather than a physiological mechanism regulating insulin translation. For these reasons, we assessed the glucose response of MIN6-K8 cells^49^, a clone of the MIN6 cells that do not respond to physiologically low glucose concentrations. Indeed, MIN6-K8 cells incubated with 2.8 mM glucose did not enhance the biosynthesis of secretory granule proteins compared to glucose starvation, while their proforms were clearly increased upon stimulation with 25 mM glucose (suppl. fig. 4d).

The stress response of the MIN6 and MIN6-K8 cells was assessed by comparing eIF2α and AMPKα phosphorylation at different glucose concentrations. The phosphorylation of eIF2α upon stress induces translational arrest, and the stalled 48S preinitiation complexes are convoyed to stress granules^50^. AMPKα phosphorylation is a measure of the energy and nutrient status of a cell: low glucose and energy status induces the phosphorylation of the α-subunit of AMPK, leading to a translational arrest through the mTOR pathway^51^. eIF2α was phosphorylated in both MIN6 cells and the MIN6-K8 cells at 0 mM glucose (suppl. fig. 4e), and stress granules were present (suppl. fig. 3c and 3d). In the presence of 2.8 mM glucose, eIF2α phosphorylation was significantly reduced in both cell lines (suppl. fig. 4e). MIN6 cells also had reduced AMPKα phosphorylation at 2.8 mM glucose, which likely explains their ability to synthesize secretory granule protein proforms in these conditions. In contrast, the MIN6-K8 cells had reduced AMPKα phosphorylation only in conditions of 25 mM glucose (suppl. fig. 4e), which enabled the glucose-induced biosynthesis of secretory granule proteins (suppl. fig. 4d). The combination of the eIF2α dephosphorylation and AMPKα phosphorylation patterns in resting MIN6-K8 cells suggests that their response to a physiological low glucose concentration (2.8 mM) reflects their nutrient and energy status rather than cell stress, similarly to healthy islet beta cells.

Certain types of stress granules form even in the absence of eIF2α phosphorylation: hyperphosphorylation of 4E-binding proteins by mTOR disrupts the eIF4E-4G interaction^52^ and leads to the stalling of the preinitiation complex, which initiates stress granule formation^53^. We discovered that MIN6-K8 cells rested at 2.8 mM glucose, G3BP1 and *Ins1* mRNA colocalized in stress granules, which we identified as part of the insulin mRNP (fig. 1d and 1f, and suppl. figure 1d). The G3BP1^+^ and *Ins1^+^* mRNA stress granules dissolved upon glucose stimulation (fig. 5a). In MIN6-K8 line, hnRNP A2/B1 was dispersed throughout the whole cell volume regardless of glucose concentration (fig. 5a). We verified that hnRNP A2/B1, PTBP1, hnRNP K and DDX1, along with some of the novel RBPs of the 5’-UTRs of mRNA for secretory granule proteins also bind these sequences in the MIN6 K8 cells (suppl. fig. 5a).

**Figure 5.**
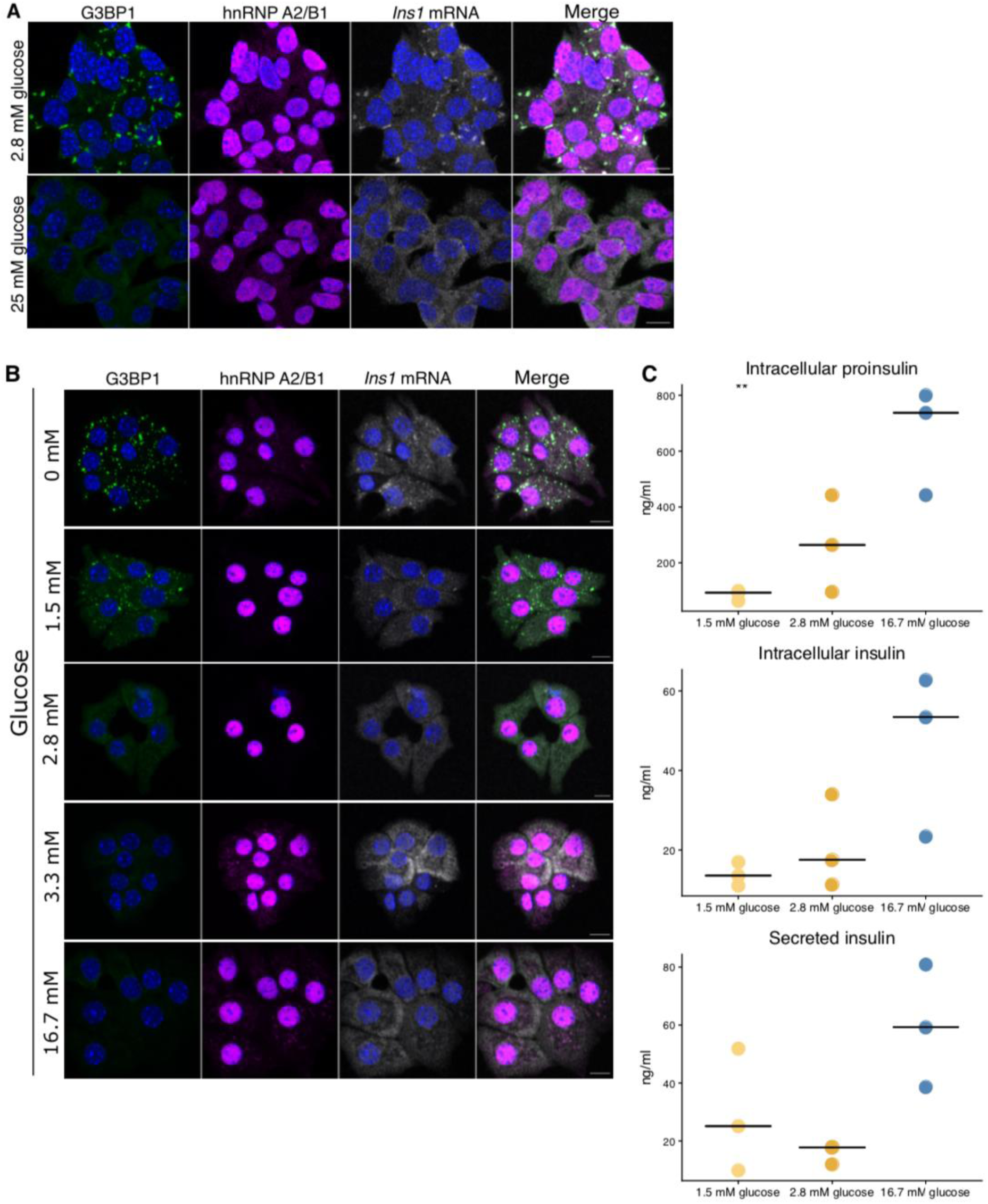
The insulin mRNA colocalizes with stress granules in resting MIN6-K8 cells. **A:** Resting MIN6-K8 cells contain G3BP1^+^ and *Ins1* mRNA^+^ stress granules that were dispersed upon glucose stimulation. Nuclei were counterstained with DAPI, maximum intensity projections are shown. Scale bar = 10 µm. R – resting, 2 h, 0 mM glucose; S – stimulated, 25 mM glucose, 2h. Ins1 – insulin1. **B:** G3BP1 and *Ins1* mRNA localized to stress granules in dispersed mouse islets incubated at 0 mM and 1.5 mM glucose and are dispersed in the cytoplasm at higher glucose concentrations. **C:** ELISA of intracellular proinsulin, insulin and secreted insulin from dispersed mouse islets incubated at different glucose concentrations shows reduced proinsulin content at 1.5 mM glucose. ANOVA: ** = p<0.01; n = 3, horizontal lines show median value.

### RNA granules in primary beta cells

Since MIN6-K8 cells rested in 2.8 mM glucose displayed G3BP1^+^ stress granules, we investigated the presence of these structures in primary mouse islet cells. The threshold for glucose-stimulated insulin biosynthesis is lower than for insulin secretions, which ensures that beta cells always have reserve stores of insulin secretory granules^3, 54^. In dispersed mouse islet cells, *Ins1/2* mRNA^+^ and G3BP1^+^ stress granules were present upon incubation with 0 mM and 1.5 mM glucose, and at higher glucose concentrations were distributed throughout the whole cytoplasm (fig. 5b). Dispersed mouse islets incubated at 1.5 mM glucose had reduced proinsulin content compared to 2.8 mM and 16.7 mM glucose (fig. 5c), indicating that insulin biosynthesis in mouse islets is already activated at 2.8 mM, hence the absence of stress granules.

Finally, we investigated the presence of stress granules in primary human beta cells. To this aim we exploited pancreas sections of metabolically phenotyped patients who had undergone pancreatectomy^55, 56^ and co-immunostained them for hnRNP A2/B1 and insulin or glucagon (fig. 6a and suppl. fig. 6a). Clinical parameters of living donors are shown in Table 1. hnRNP A2/B1^+^ cytosolic granules were present in the beta cells of three ND and two T2D patients (arrows in fig. 6 and suppl. fig. 6), although in most of the cells hnRNP A2/B1 was restricted to the nucleus, as in the bottom panel of suppl. fig. 6a. A small number of the hnRNP A2/B1^+^ structures colocalized with G3BP1^+^ structures in the sections (suppl. fig. 6b). Using RNAscope, we found that *INS* mRNA partially resides in structures similar in size, shape and localization to the hnRNP A2/B1^+^ granules (fig. 6b and suppl. fig. 6c). While in beta cells of some patients *INS* mRNA presented a dispersed pattern (fig. 6b), in most cases we could detect *INS* mRNA^+^ structures which resembled hnRNP A2/B1^+^ granules. The occurrence of hnRNP A2/B1^+^ and *INS* mRNA^+^ structures did not depend on the diabetes status of the subjects. Co-staining of *INS* mRNA with hnRNP A2/B1 and stress granule markers was not possible due to technical limitations^57^. Nonetheless, these findings provide the first evidence for the enrichment of *INS* mRNA in RNA granules in human beta cells, and further research is needed to define their identity and function in health and disease.

**Figure 6.**
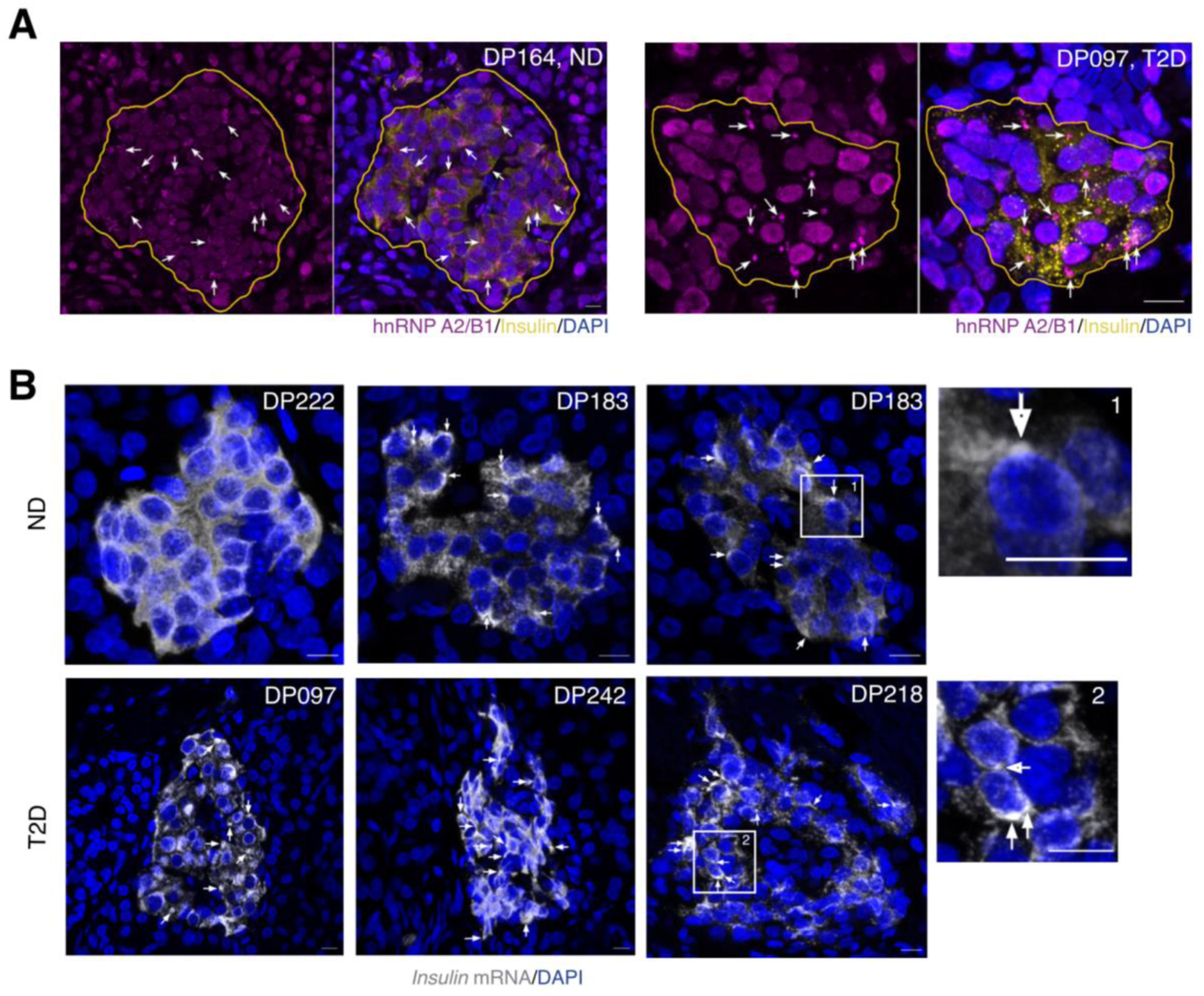
hnRNP A2/B1 and *insulin* mRNA-positive stress granules in primary beta cells. **A:** hnRNP A2/B1^+^ stress granules (arrows) were detected in FFPE pancreatic sections from partially pancreatomized ND and T2D patients. Insulin marks beta cells and islets are circled with yellow lines. **B:** *INS* mRNA localized to cytoplasmic RNA granules (arrows) in FFPE pancreatic sections from partially pancreatomized ND and T2D patients. Scale bar = 10 µm. ND – non-diabetic; T2D – type 2 diabetic; FFPE – formalin fixed paraffin embedded.

**Table 1.** Clinical parameters of living pancreas donors. BMI – body mass index; M – male; F – female; ND – non-diabetic, T2D – type 2 diabetic; hba1c – glycated haemoglobin.

| patient id<br>[DP-Nr] | diabetes<br>status | gender | age | bmi | hba1c | glucose<br>(mmol/l)<br>OGTT<br>120min | HOMA1<br>IR | duration<br>of known<br>diabetes<br>[years] | oral antidiabetics | insulin |
| --- | --- | --- | --- | --- | --- | --- | --- | --- | --- | --- |
| DP159 | ND | M | 52 | 26.80 | 4.8 | 3.02 | 0.30 | - | none | none |
| DP173 | ND | F | 61 | 19.43 | 5.3 | 5.58 | 3.17 | - | none | none |
| DP183 | ND | M | 60 | 23.31 | 4.9 | 5.95 | 3.86 | - | none | none |
| DP222 | ND | F | 65 | 26.03 | 5.2 | 5.44 | 2.34 | - | none | none |
| DP164 | not<br>assignable | F | 86 | 30.40 | 5.0 | NA | no calc | 0 | none | none |
| DP097 | T2D | M | 30 | 25.98 | 7.7 | NA | 0.68 | 3 | none | Lantus 16-0-0-0IE,<br>NovoRapid 26-4-10IE |
| DP099 | T2D | F | 66 | 29.71 | 8.4 | NA | 23.23 | 11 | none | Humaninsulin 14-14-14 IE,<br>Basalinsulin 0-0-0-16IE |
| DP166 | T2D | F | 76 | 37.92 | 7.1 | NA | 5.80 | 23 | Januvia 100mg<br>1-0-0-0 | Insulin lispro Humalog 26-24-26-0 I.E.,<br>Insulin glargin Lantus 0-0-0-26 I.E. |
| DP201 | T2D | M | 70 | 26.83 | 6.4 | NA | 3.30 | 11 | Siofor 850 1-0-1,<br>Onglyza 5mg 1-0-0 | none |
| DP218 | T2D | F | 76 | 28.89 | 6.1 | NA | 10.23 | 30 | none | Huminsulin Basal 0-0-0-18 I.E.,<br>Huminsulin Normal 16-12-12-0 I.E. |
| DP242 | T2D | M | 74 | 25.30 | 8.1 | NA | 4.60 | 1 | Metformin 1000mg<br>1/2-1/2-1-0 | none |

## Discussion

Pancreatic beta cells use post-transcriptional mechanisms to regulate insulin and secretory granule biosynthesis, ensuring that new insulin granules can quickly replace the secreted ones. A number of proteins regulate insulin expression^6, 35, 37, 58^ among which PTBP1 and hnRNP K have been suggested to coordinate the biosynthesis of insulin and several other secretory granule proteins^16, 29, 30^. However, a comprehensive overview of the secretory granule protein mRNPs has been missing. In this study, we determine that under resting conditions, one set of RBPs binds to secretory granule protein mRNAs, which target them for storage in stress granules. Glucose induces RNP remodelling, causing the binding of another set of RBPs to the transcripts, the dissolution of stress granules and a burst in secretory granule protein biosynthesis. Notably, stress granules are also present in human beta cells. Moreover, we determined that hnRNP A2/B1 is a novel factor for control of *Insulin1* mRNA translation.

To our knowledge this is the first study to use an unbiased approach to compare glucose - induced changes in protein binding to mRNAs of insulin secretory granule proteins with the 5’-UTR of a housekeeping mRNA. As such, our findings significantly contribute to the understanding of glucose stimulated post-transcriptional regulation in beta cells. Using *in vitro* RNA pull-downs, we discovered glucose-dependent differences in RBP binding, which could be explained by increased expression of the corresponding RBPs, their nucleocytoplasmic shuttling or glucose-induced post-translational modifications. Among the proteins that were identified by RNA pull-down, we found some of the proteins previously known to regulate insulin expression. In particular, previous studies have shown that mRNAs for secretory granule proteins are co-regulated by PTBP1^16, 30^ and hnRNP K (our unpublished results). We found that besides PTBP1, six novel RBPs specifically co-bind to the transcripts for secretory granule proteins in resting conditions. We also discovered novel RBPs for these transcripts in stimulated conditions. DDX1, which binds to the insulin mRNA in rat insulinoma INS-1 cells^38^ also interacts with all of the secretory granule protein transcripts that we tested. Additionally, several of the identified proteins have been associated with neuronal granules^59, 60^ and stress granules^61^.

In contrast to previous studies, however, we did not detect significant enrichment in the binding of TIAR, HuD, PABP1 or PDI to any of the tested transcripts in MIN6 cells. While all of these studies used a similar methodology for the RNA pull-downs, the conflicting results can be explained by differences in the model system (cell lines vs. islets), the starting material (cytosolic vs. whole cell extracts), the RNA sequence used for pull-down (full length 5’-UTR vs. oligonucleotide segments) and in the identification method (WB vs. MS). Most importantly, each of the previous studies analyzed specific binding to the 5’-UTR of insulin transcripts using different control sequences: a scrambled sequence with the same nucleic acid composition^37^, a deletion fragment of rat *Ins1* 5’-UTR^58^, or the 3’-UTR of *Gapdh* mRNA^35^. While such controls can account for unspecific RNA binding, they are not appropriate for testing how beta cells selectively up-regulate the expression of secretory granule proteins compared to the rest of the cellular proteome. To this aim, we used the full length 5’-UTR of a non-secretory granule protein mRNA to determine the RBPs that differentially regulate secretory granule protein expression compared with other beta cell genes.

Nevertheless, our approach also has several limitations which do not exclude false positive and negative results. First of all, while our assays were designed in a way that limits unspecific binding, the RNA pull-downs were still performed *in vitro*. Therefore, we cannot rule out the formation of non-functional RNA-protein complexes due to post-lysis effects. Additionally, since we previously showed that secretory granule protein mRNAs undergo cap-independent translation, we performed our pull-downs using sequences that did not have a 5’-cap. Accordingly, some proteins binding to the 5’-UTR of *γ-tubulin*, which is translated in a cap-dependent fashion, might have been missed. Moreover, the UV crosslinking, which was used to stabilize the RNA-protein interactions by forming covalent bonds, could enable the retention of unspecific proteins, even after stringent washing. Furthermore, UV cross-linking only links direct RNA-protein contact and may not capture the larger multi-protein complexes that make up the overall RNP architecture *in vivo*. Finally, since we controlled for unspecific binding using the 5’-UTR of a housekeeping mRNA, it is possible that RBPs which specifically bind and regulate both control and target mRNAs would be classified as background. In the future, the integration of aptamer sequences, e.g., the S1 streptavidin binding aptamer, into the target transcripts could enable the purification of *in vivo* crosslinked RNA-protein complexes and overcome the limitations of this study.

A novel interacting partner identified in our screen is hnRNP A2/B1, which is one of the seven enriched RBPs shared among the mRNAs for secretory granule proteins in resting conditions. We discovered that the 5’-UTRs of transcripts for secretory granule proteins contain putative A2REs and verified that the binding of hnRNP A2/B1 decreases upon the mutation of these elements in the *Ins1* mRNA 5’-UTR. The A2RE motif does not overlap with the exon-exon junction site in the *Ins1* mRNA 5’-UTR, suggesting that hnRNP A2/B1 is not involved in splicing within the 5’-UTR of this transcript. The mutation of the A2REs reduced the translational efficiency of a downstream luciferase reporter, although there were no differences in luciferase intensity between resting and stimulated cells, which confirms previous results that both the 5’- and 3’-UTR cooperate to increase glucose stimulated insulin biosynthesis^9^.

In mouse beta cells most insulin results from the translation of *Ins2* mRNA^40^. In our MIN6 cells *Ins2* mRNA is twice as abundant as *Ins1* mRNA. In *Hnrnpa2b1* KO MIN6 cells the insulin protein content was significantly reduced, although only the levels of *Ins1* mRNA were decreased, while those of *Ins2* mRNA were unchanged. Glucose stimulation upregulated insulin biosynthesis in both *WT* and *Hnrnpa2b1* KO cells, as indicated by the increased amount of intracellular proinsulin. However, proinsulin levels in *Hnrnpa2b1* KO cells were significantly lower. Hence, it is conceivable that hnRNP A2/B1 regulates *Ins1* mRNA stability and *Ins2* mRNA translation and that a combination of both mechanisms accounts for reduction in insulin protein content in its absence. Interestingly, the nucleotide sequences of the *Ins1 and Ins2 mRNAs* differ mainly in their 5’-UTRs.

RNA pull-down assays indicated the binding of hnRNP A2/B1 to *Pc2* and *Ica512* mRNA 5’-UTRs, which contain putative A2REs. However, its deletion affected neither mRNA nor protein levels of PC2 and ICA512. Additionally, glucose stimulation increased the proforms of these proteins to a similar extent in both WT and *Hnrnpa2/b1* KO cells. Thus, it seems that beta cells rely on a fail-safe mechanism to support secretory granule production even in the absence of hnRNP A2/B1. Consistent with this hypothesis, we found that hnRNP A1 levels in *Hnrnpa2/b1* KO MIN6 cells were increased compared with wild-type cells. hnRNP A2/B1 is structurally and functionally similar to its paralogue hnRNP A1, which was also among the common enriched RBPs for the 5’-UTRs of secretory granule protein mRNAs. Possibly increased hnRNP A1 expression can compensate for the loss of hnRNP A2/B1 and enable beta cells to sustain the biosynthesis of PC2, PC1/3 and ICA512, albeit not that insulin.

Interestingly, we discovered that hnRNP A2/B1 and the *Ins1* mRNA localize to stress granules in the cytoplasm of resting insulinoma cells, which dissolve upon glucose stimulation. While the presence of stress granules has been previously observed in thapsigargin-treated rat insulinoma INS-1 cells^37^, to our knowledge, this is the first time RNA granules are reported to have a physiological function in beta cells. Although MIN6 and MIN6-K8 cells differ in the presence of hnRNP A2/B1 in stress granules, G3BP1^+^ RNA granules were present in both cell types at their respective “low glucose” concentrations and colocalized with *Ins1* mRNA. Notably, hnRNP A2/B1, eIF3b and G3BP1 localize to stress granules, but they have also been linked to RNA granules independent of a stress stimulus. For instance, in neurons and oligodendrocytes, hnRNP A2/B1 regulates the cytoplasmic transport of a set of mRNAs by sequestering them into RNA granules. Once these reach their final destination, hnRNP A2/B1 fosters translation in response to stimuli (i.e. synaptic transmission)^44, 48, 62^.

In surgical samples of metabolically phenotyped pancreactomized patients with normoglycemia or T2D, we detected hnRNP A2/B1^+^ and *INS^+^* RNA granules in the cytoplasm of human beta cells. These data provide first evidence for the presence of RNA granules in primary human beta cells. Due to technical limitations, we were unable to simultaneously label FFPE sections for hnRNP A2/B1 and *INS* mRNA. Intriguingly, we could not reveal differences for the presence of RNA granules in beta cells of subjects with T2D relative to normoglycemic donors. Since these patients were fasted overnight prior to surgery, and may also be differently treated with glucose and/or insulin during the prolonged surgical procedure, we cannot draw conclusions regarding possible differences about RNA granules in relation to glucose tolerance. However, given the lower levels of G3BP1 mRNA in islets of donors with T2D^63^, this scenario deserves to be considered further.

In general, there is emerging evidence about the role of RBPs in beta cell dysfunction. PTBP1 has been linked to beta cell dysfunction in human islets^32, 33^ and polymorphisms in *PTBP1* have been associated with reduced glucose tolerance^34^. DDX1 has been linked to the palmitate-feeding induced reduction in insulin translation in mouse islets. Neurodegenerative diseases, such amyotrophic lateral sclerosis, frontotemporal dementia and multiple sclerosis have been linked to the formation of pathological inclusions, where mutations in RBPs alter stress granule dynamics, compromising the gene expression profile and cell function^46, 64–68^. Notably, while hnRNP A2/B1 has not yet been associated with T2D, mutations in its prion-like domain can cause abnormal protein aggregation and the formation of amyloid plaques in amyotrophic lateral sclerosis^46^.

As a sens0r of nutrient status, AMPKα phosphorylation in conditions of low nutrient availability initiates stress granule formation through the mTOR-4E-BP pathway^53^. AMPK inhibition through dephosphorylation of Thr172 in the AMPKα subunit in glucose-stimulated beta cells has been well studied in mouse, rat and human islets^51^. We demonstrated this in MIN6 cells (suppl. fig. 4d and 4e). Previous studies show that AMPK inhibition in stimulated beta cells augments insulin secretion^69^, albeit AMPKα dephosporylation happens at a glucose concentration that does not initiate insulin secretion^70^, but, presumably, can initiate insulin biosynthesis^3, 54^. In models of diet induced obesity, glucose stimulation failed to inhibit AMPK activation in islets from high-fat fed mice ^71^. In human islets, diabetogenic conditions did not alter AMPK expression, but chronic fructose exposure increased p-AMPK levels to fasting levels^72^ and AMPK activation was reduced in T2D islet donors compared to ND controls^73^. Here, we link AMPKα inactivation with insulin biosynthesis. These data, in conjunction with our discovery of hnRNP A2/B1^+^ stress granules in human islets warrants further investigation into the post-transcriptional mechanism regulating insulin expression.

A hallmark of T2D is impaired beta cell function and insulin secretion. In our model, deletion of a RBP which regulates insulin biosynthesis resulted in decreased insulin secretion, even though the insulin stimulation index of WT and *Hnrnpa2b1* KO cells was comparable. Similarly, depletion of *Ptbp1* in INS1 cells reduced glucose-stimulated biosynthesis of secretory granules and their stores^30^. Also, some monogenetic forms of diabetes are caused by mutations in factors promoting translation, such as CDKAL1 and TRMT10^74^. Taken together, these data point to deficits in insulin biosynthesis as possible causes of diabetes, even though the secretory machinery of beta cells is intact. Further studies on the role of RBPs and stress granules for post-transcriptional regulation of mRNAs coding for secretory granule proteins will therefore contribute to the understanding of beta cell physiology in healthy and pathological conditions.

## Author contributions

Conceptualization: J.V., M.S; Methodology: J.V., Dj.V.; Validation: J.V.; Formal analysis: J.V., Dj.V.; Investigation: J.V., Dj.V, K.G., E.D.Q.D.; Resources: C.W., C.M., A.S., E.S., D.F, N.K., M.D. M.S.; Writing - Original Draft: J.V.; Writing - Reviewing & Editing: J.V., Dj.V., M.S.; Visualization: J.V.; Supervision: M.S.; Funding Acquisition: J.V., M.S.

## Acknowledgments

We would like to thank Martin Neukam for the detailed reading of the manuscript and optimizing islet dispersion, Katja Pfriem for administrative assistance and members of the Solimena lab for their feedback and discussion. The work in the Soliman lab was supported by the German Center for Diabetes Research (DZD e.V.), which is financed by the German Ministry for Education and Research.

## Materials and Methods

### Cell culture and islet culture

Subclones of the MIN6 mouse insulinoma cell line were cultured as previously described^30^. For stimulation, cells were first pre-incubated in resting medium (15 mM HEPES, pH 7.4, 5 mM KCl, 350 mM NaCl, 24 mM NaHCO_3_, 1 mM MgCl_2_, 2 mM CaCl_2_, 0 mM/2.8 mM glucose, 1 mg/ml ovalbumin) for 1 h, followed by resting or stimulation medium (15 mM HEPES, pH 7.4, 25 mM glucose, 5 mM KCl, 120 mM NaCl, 24 mM NaHCO_3_, 1 mM MgCl_2_, 2 mM CaCl_2_, 25 mM glucose, 1 mg/ml ovalbumin) for 2 h. Transfections were performed using the G16 program of the AmaxaTM Cell Line NucleofectorTM Kit V (Lonza AG), following the manufacturer’s instructions.

Pancreatic islets were isolated from C57BL/6 mice, dispersed and cultured as previously described^75^ with minor adjustments, namely the use of accutase instead of trypsin and the coverslips being coated with ECM (Sigma, E1270). The dispersed islets were incubated in resting medium with the appropriate glucose concentration for 2 h.

### Cell extract preparation

Cells were lysed in lysis buffer (20 mM Tris-HCl pH 8.0, 140 mM NaCl, 1 mM EDTA, 1% v/v Triton X-100, 1X protease inhibitor cocktail P8340 (Sigma-Aldrich Chemical Co.), 1X phosphatase inhibitor cocktail (Merck Chemicals), unless otherwise stated. Nuclear and cytosolic extracts were prepared using the NE-PER™ Nuclear and Cytoplasmic Extraction Reagents kit (Thermo Scientific).

### *In vitro* transcription and biotinylation

The 5’-UTRs of transcripts were amplified from MIN6 cell cDNA with the addition of a T7 promoter and *in vitro* transcribed using the AmpliScribe™ T7-Flash™ RNA transcription kit (Ambion) and biotin-16-CTP (Roche) in accordance with the manufacturer’s instructions. The RNA products were visualized by denaturing RNA gel electrophoresis.

### RNA pull-down

400 pmol of biotinylated RNA was bound to 100 µl Dynabeads® M-280 Strepavidin (Invitrogen) in RNA-binding buffer (20 mM HEPES pH 7.4, 50 mM EDTA, 5% glycerol, 50 mM KCl, 3 mM MgCl, 1 mM DTT, 20U SUPERase RNase inhibitor (Invitrogen)) rotating at 4 °C for 1 h. Cytosolic protein lysates from MIN6 cells were precleared with 20 µg yeast tRNA (Ambion) and 50 µl Dynabeads by rotating at 4°C for 1 h. The RNA-bound beads were washed twice in RNA-binding buffer and incubated with 125 - 800 µg of precleared protein extracts, in a total volume of 500 µl protein binding buffer (20 mM HEPES pH 7.4, 50 mM EDTA, 5% glycerol, 50 mM KCl, 3 mM MgCl, 1 mM DTT, protease inhibitors 1:100, phosphatase inhibitors 1:100, 20U RNase inhibitor, 20 µg yeast tRNA), as triplicates. The binding was performed at 4 °C for 2 h with rotation. Protein-RNA interactions were stabilized by UV irradiation for 30 min in the UVC 500 UV crosslinker (Amersham), after which the beads were washed three times in 500 µl protein binding buffer with 0.05% Triton X-100 and two times in RNA-binding buffer. Proteins were eluted by digesting the RNA with 3 µg RNase A (Roche) and 30U RNase T1 (Roche) in 400 µl elution buffer (20 mM HEPES pH 7.4, 50 mM EDTA, 5% glycerol, 50 mM KCl, 3 mM MgCl, 1 mM DTT, protease inhibitors 1:100), overnight at 4 °C. Eluates were precipitated overnight with 100% ethanol. Pellets were dissolved in 1X SDS buffer and loaded on SDS-PAGE or prepared for MS analysis.

### RNA-IP

MIN6 cells were grown in 10 cm dishes until 80% confluent and UV crosslinked for 10 min in the UVC 500 UV crosslinker (Amersham). Cell pellets were dissolved in RNA lysis buffer (50 mM Tris-HCl, pH 7.5, 100 mM NaCl, 6mM MgCl_2,_ 1% NP-40, 0.1 % SDS, 0.5% sodium deoxycholate, 1:100 protease inhibitors, 20 U/ml SUPERase RNase inhibitor) and homogenized by passing through a 20G needle. The lysates were incubated for 15 min on ice and treated with DNase I for 15 min at room temperature and centrifuged at full speed. Supernatants were precleared with Protein G Dynabeads and 5 µg mouse IgG, rotating at 4 °C for 1 h, after which protein concentration was measured and 1/10 of the sample was removed for total RNA and protein input. IPs were performed in IP buffer (50 mM Tris, 100 mM NaCl, 0.05% NP-40, 20U SUPERase RNase inhibitor, 1:100 protease inhibitors) with 500 µg protein and 10 µg of anti-hnRNP A2/B1 from Santa Cruz Biotechnology Inc. (sc-374053) or mouse IgG antibody from Jackson ImmunoResearch Lab (015-000-003). The samples were incubated for 2 h, rotating at 4 °C, after which 100 µl Protein G Dynabeads were added and incubated for an additional hour. The samples were washed twice in high-salt buffer (50 mM Tris-HCl, pH 7.5, 1 M NaCl, 1mM EDTA, 0.1% SDS, 1% NP-40) and twice in RNA lysis buffer, rotating for 2 min for each wash. The beads were resuspended in 500 µl proteinase K buffer (5 mM Tris-HCl, pH 7.5, 1 mM EDTA, 0.05 % SDS, 20 U SUPERase RNase inhibitor) and 1/5 of the sample was taken to check the IP by Western blot. 4.2 µl of proteinase K (20 mg/ml stock) was added and the samples were incubated for 90 min at 50 °C. RNA was extracted by phenol/chloroform and precipitated with ethanol. Pellets were resuspended in 12 µl DEPC water, 2 µl of which was used for reverse transcription and subsequent qPCR.

### CRISPR/Cas9 knock-out

Guide RNA (gRNA) sequences specifically targeting the mouse *Hnrnpa2b1* gene sequence were designed using the online CRISPR design tool available at http://crispr.mit.edu/. A 20 nt long high-quality guide sequence with a low number of potential off-target effects was chosen based on the proximity to the Cas9 cleavage site to the start codon. Complementary oligos were designed with overhangs that enable ligation into the BbsI-digested pSpCas9(BB)-2A-Puro plasmid. The oligo sequences were: ACCGAGTCCGCGATGGAGGTAAC and AAACGTTACCTCCATCGCGGACTC. Selection of transfected cells was achieved by adding puromycin to the culture medium, which was maintained in the media for further culturing. Single cells were plated by limited dilution and expanded until confluent. Clones were screened for hnRNP A2/B1 expression by Western blot and editing was confirmed with sequencing.

### Western blot

Non-denaturing TRIS-Tricine gels were used for visualization of insulin, all other proteins were separated and immunoblotted using standard SDS-PAGE and Western blot techniques. The following primary antibodies were used: anti-PTBP1 from Invitrogen Corp. (32-4800), anti-hnRNP K from Santa Cruz Biotechnology Inc. (sc-28380), anti-hnRNP A2/B1 from Santa Cruz Biotechnology Inc. (sc-374053), anti-hnRNP A2/B1 from Sigma Aldrich Chemical Co. (R4653), anti-G3BP1 from abcam (ab181150), anti-Insulin from Sigma-Aldrich Chemical Co. (I-2018), anti-p-eIF2α from Signaling Technology (#9721), anti-eIF2α from Signaling Technology (#9722), anti-PC1/3 from GeneTex (GTX113797), anti-ICA512 (self-raised) anti-p-AMPK from Cell Signaling Technology (#2531), anti-AMPK from Cell Signaling Technology (#2532), anti-DDX1 from LS Bio (LS-C334806), anti-hnRNP Q from Proteintech (14024-1-AP), anti-Fxr1 from Thermo Fisher (702410), anti-RPS6 from (#2212) Cell Signaling Technology, anti-hnRNP A1 from Sigma-Aldrich Chemical Co. (R4528), anti-PC2 from GeneTex (GTX114625), anti-gamma-tubulin from Sigma (T-6557). Secondary HRP-conjugated antibodies were from Bio-Rad, except for IP samples, where anti-mouse light chain specific HRP-conjugated antibodies (Dianova) were used to avoid interference from the heavy chain of the precipitated IgG. Nitrocellulose membranes were developed with SuperSignal West Pico Chemiluminescent Substrate or SuperSignal West Femto Maximum Sensitivity Substrate (Thermo Fisher Scientific) according to the manufacturer’s instructions. Bands were detected using the Amersham Imager 600 (GE Healthcare). Band intensity was quantified using ImageStudioLite (Li-COR) and normalized to that of the loading control (γ-tubulin).

### ELISA

After glucose stimulation, media was collected and the cells were washed twice in PBS, scraped in acid ethanol. All samples were stored at –20 °C until use. Insulin and proinsulin were measured using the Insulin ELISA kit (ALPCO) and proinsulin ELISA kit, respectively, according to the manufacturer’s instructions. DNA was precipitated from the acid ethanol samples and used for normalization.

### RNA isolation and qPCR

Total RNA was isolated using the RNeasy Mini Kit (Qiagen Inc.) including the on-column DNase digestion step with the RNase-Free DNase Set (Qiagen Inc.). One µg RNA was reverse transcribed using the MLV RT kit (Promega). The GoTaq^®^ qPCR Master Mix (Promega) was used according to manufacturer’s instructions and the samples were run in triplicate on the AriaMx Real-time PCR System (Agilent). Relative quantification was performed using the 2^-ddct^ method. Reference gene selection was achieved by testing the expression stability of the following housekeeping genes: *β-actin*, *γ-tubulin*, *β2-microglobulin*, *HPRT* and *TBP*. The most stable reference gene was determined for each experiment using the RefFinder^76^.

### Immunofluorescence

For immunofluorescence, cells were grown on coverslips using standard culture conditions, fixed with 4% PFA for 20 min at room temperature and permeabilized in 0.1% Triton X-100 in PBS for 10 min at room temperature. Coverslips were incubated with blocking buffer (0.5% BSA, 0.2% fish skin gelatine solution in PBS, filter sterilized) for 30 min, primary antibodies diluted in blocking buffer for 30 min, and AlexaFlour-coupled secondary antibodies (1:200) from the appropriate host species. Double stainings were performed with primary antibodies from different host species; nuclei were counterstained with DAPI. Coverslips were mounted on glass slides using Mowiol (24% w/v glycerol, 9.6% w/v mowiol, 96 mM Tris-HCl pH 8.5, 2.5% DABCO). Primary antibodies used were: anti-hnRNP A2/B1 from Santa Cruz Biotechnology Inc. (sc-374053), anti-G3BP1 from abcam (ab181150), anti-Insulin from Sigma-Aldrich Chemical Co. (I-2018), anti-eIF3b from Santa Cruz Biotechnology Inc. (sc-137214), and anti-Dcp1a from abcam (ab183709). For RNase treatment prior to IF and FISH, the cells were treated with 0.2 mg/ml RNase A in 10 mM Tris-HCl, pH 7.5, 15 mM NaCl, 0.05% saponin for 15 min, prior to fixation.

### Fluorescent *in situ* hybridization (FISH)

FISH probes were designed using the Stellaris Probe Designer tool. The probes were BLASTed to check specificity: if more than 5 probes bound to a non-target gene with 16 nt or more, they were removed from the probe set. The mouse *gapdh* probe set from Stellaris was used as a positive control. Cells were grown on coverslips, fixed in 4% PFA and permeabilized with 0.1% Triton X-100. Probe hybridization was performed according to the manufacturer’s instructions. When combining FISH with IF, the IF was performed first, after which the cells were fixed for 10 min in 4% PFA and hybridized to FISH probes overnight.

### Confocal microscopy

Images were acquired with a confocal laser scanning Zeiss LSM 780 microscope (Carl Zeiss AG) equipped with a 63x/1.46 Oil DIC (Plan-Apochromat) objective. Image acquisition was done in the sequential scanning mode with a line averaging of 2. Optical sections were taken throughout the whole z-volume of the cells, with a z-step size set to half of the optical section thickness. Microscopy images were processed using Fiji, maximum intensity projections are shown. Background fluorescence intensity was measured using an ROI and the average intensity value was subtracted from the full image. If modified, the brightness and contrast settings were adjusted for the whole image, with identical settings for all images within one experiment. Colocalization analysis was performed on background subtracted maximum intensity projections using the Coloc2 Fiji plugin.

### Luciferase assays

The Rosa26 promoter and the 5’UTRs of secretory granule protein transcripts were subcloned into the pNL1.1 (Promega) using standard cloning techniques. Mutagenesis of the A2REs in the *Ins1* 5’-UTR was performed with the QuikChange II Site-directed mutagenesis kit (Agilent). Cells were co-transfected with each pNL1.1 construct and the pGL4.54 for normalization. Cells were lysed in Glo Lysis Buffer (Promega) four days after transfection and luciferase activity was measured using the Nano-Glo® Dual-Luciferase® Reporter Assay System using the Synergy Neo2 multi-mode reader (BioTek). Nanoluciferase activity was normalized to that of the firefly luciferase.

### Staining of human pancreas sections

Surgical pancreas tissue from metabolically profiled, partially pancreatectomized living human donors used in this study were obtained from the IMIDIA biobank and have been previously described^55^. 5 µm FFPE sections were mounted on microscope slides and prepared using standard procedures; antigen retrieval was performed by boiling the samples in citrate buffer with 1% Tween20 for 3 min. Sections were blocked using Dako Antibody dilutent. Primary and secondary antibody incubation was performed in Dako Antibody dilutent for 1 hour. Nuclei were counterstained with DAPI and coverslips mounted with Mowiol. anti-hnRNP A2/B1 from Santa Cruz Biotechnology Inc. (sc-374053), anti-G3BP1 from Proteintech (13057-2-AP), anti-glucagon from Dako (A0565), Anti-Insulin Alexa Fluor® 488 from Invitrogen (53-9769-82). For FISH, *INS* mRNA specific probes were used with the RNAscope^®^ Multiplex Fluorescent Assay Kit (Advanced Cell Diagnostics), according to the manufacturer’s instructions.

### Sample prep for MS

Pellets from RNA pull-downs were solubilized in 50 μl denaturation buffer (6 M urea, 2 M thiourea, 50 mM ammonium bicarbonate, pH 8.5) and reduced by addition of 2 μl DTT (10 mM) for 30 min at room temperature. Cysteines were alkylated using 55 mM iodoacetamide and incubated in the dark for 20 min at room temperature. The samples were pre-digested with 1µg LysC endopeptidase for 3 h at room temperature, after which they were diluted four times with 50 mM ammonium bicarbonate, pH 8.5 and digested with 1 µg trypsin, overnight at room temperature. The reaction was stopped by acidification of the sample to pH < 2.5 with 10% trifluoroacetic acid. The peptides were desalted and concentrated on a StageTip (Stop and go extraction Tip) containing C18 reverse-phase material (Empore, 3M, Minneapolis, USA). For LC-MS measurement, peptides were eluted from StageTips by forcing 60 µl of buffer B (80% acetonitrile, 0.1% formic acid) directly into a 96-well microtiter plate. Eluted peptide solutions were dried in a speed vac to a volume of 3-4 µl and 5 µl of buffer A (5% acetonitrile, 0.1% formic acid in water was added to each sample.

### LC-MS acquisition and post-acquisition workflow

Peptides from each sample were separated by reverse phase liquid chromatography on a 20 cm fritless silica microcolumn (inner diameter 75 µm) packed in-house with ReproSil-Pur C18 3 µm resin (Dr. Maisch GmbH, Ammerbuch, Germany) using the Proxeon 1000 system (Thermo Scientific). Peptides were separated on an 8-50% acetonitrile gradient with 0.1% formic acid, during 155 min with a flow rate of 250 nl/min. Eluting peptides were directly ionized by electrospray ionization at 2.2 kV and transferred into a Q Exactive Plus Quadrupole-Orbitrap hybrid mass spectrometer (Thermo Scientific).

Mass spectrometry was conducted in a data-dependent mode with full scan (MS1) followed by 10 fragmentation scans (MS2) of the 10 most intense ions. MS1 scans were performed in a 300-1,700 m/z range at the resolution of 70,000 and target value of 3×10^6^. MS/MS scans were performed with higher energy collision (HCD), normalized energy 26%, isolation width +/- 4 m/z, target value 2×10^5^, maximum injection time 120 ms and resolution of 35,000. Dynamic exclusion was set to 30 s and unassigned or +1 charge states were rejected for isolation.

Raw data were processed using MaxQuant software (version 1.5.1.2)^77^ with the built-in Andromeda search engine. Search was performed against the mouse proteome database (Uniprot, 2014), using the target-decoy approach. Reversed peptide sequences with K and R as special amino acids was chosen as the decoy strategy. By default, common contaminants were included in the search. Carbamidomethylation of cysteine was set as a fixed modification, while oxidation of methionine and acetylation of protein N-terminus were set as variable modifications. Digestion mode was set to Trypsin/P specific, minimum peptide length was 7 amino acids and maximum 2 missed cleavage sites were allowed. False discovery rate (FDR) was set to 0.01 for both peptide and protein identification. The “match between runs” option was chosen for transferring MS/MS identifications with the maximal retention time tolerance of 2 min. Protein quantification was performed using the label-free quantification (LFQ) algorithm^78^, where a minimum of 2 LFQ ratio counts was required. Raw data have been deposited to the ProteomeXchange Consortium via the PRIDE partner repository with the dataset identifier PXD026956

The Perseus software^41^ was used for statistical analysis and graphical representation of proteomics data. The proteinGroups.txt-file is loaded into Perseus with LFQ intensities marked as “expression.” Proteins were filtered for “contaminants”, “reverse” and “identified by site” and proteins that were not detected in at least two replicates of at least one group were deleted. Values were log2 transformed and the missing values were imputed separately for each column from a normal distribution with a width of 0.3 and downshift of 1.8. LFQ intensities were statistically analysed using a permutation-based t-test with an FDR of 5% and slope of 0.1. Volcano plots were created with the “-log t-test p-value” versus the “t-test difference” (fold change).

## Supplementary figure and table legends

**Supplementary figure S1.**
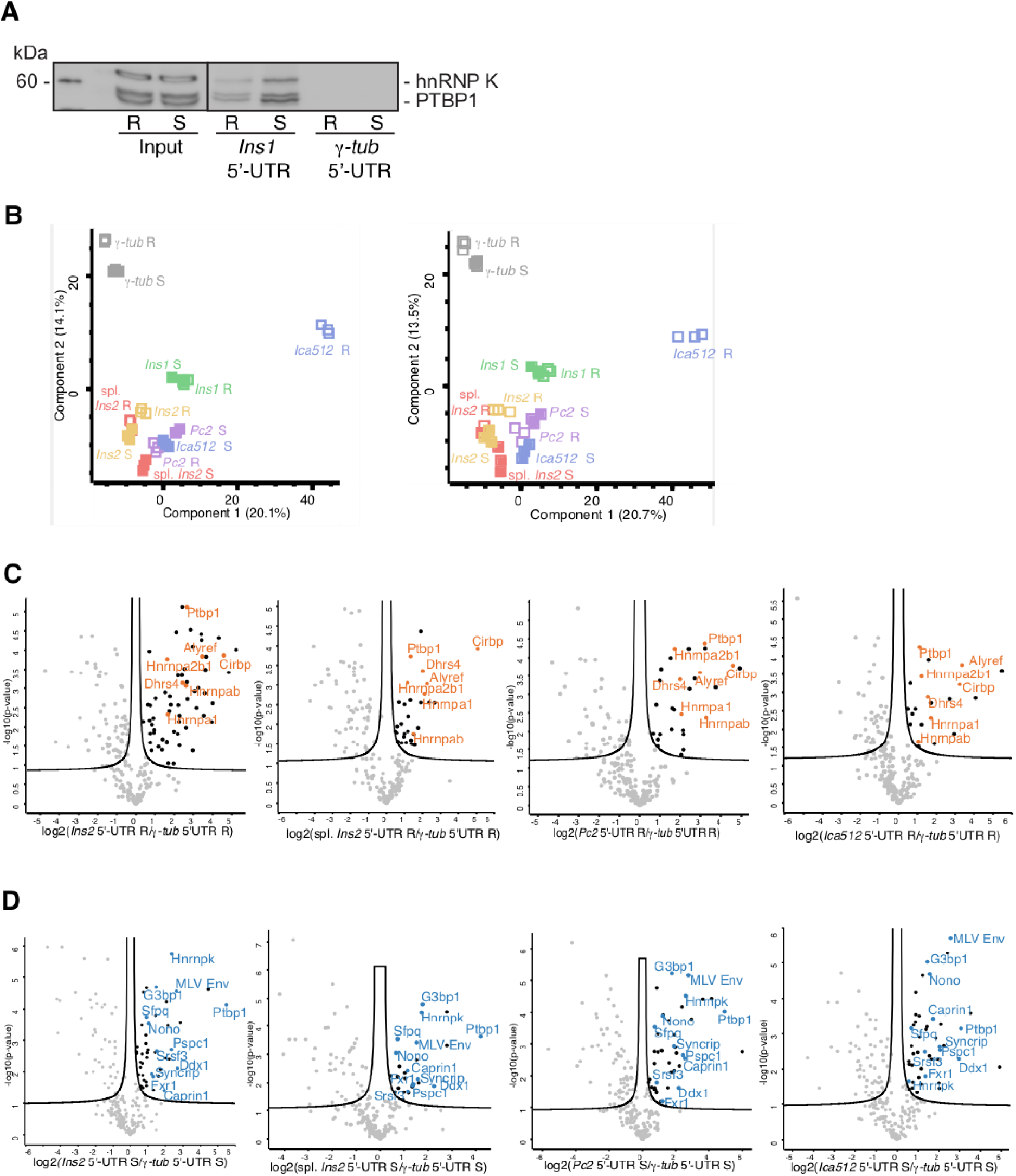
*In vitro* RNA pull-down with the 5’-UTRs of secretory granule protein transcripts. Relates to figure 1. **A:** PTBP1 and hnRNP K do not bind to *γ-tub* 5’-UTR control, validating this sequence as a non-specific control for the identification of secretory granule protein RNPs. **B:** PCA plot of two independent experiments in resting (R, open squares) and stimulated (S, filled squares) MIN6 cells. **C and D:** volcano plots showing significantly enriched proteins binding to the 5’UTRs of *Ins2,* spliced *Ins2, Pc2* and *Ica512* 5’-UTRs compared to the *γ-tub* 5’-UTR in resting (orange, **C**) and stimulated (blue, **D**) MIN6 cells. Ins – insulin; Pc2 – prohormone convertase 2; Ica512 – islet cell autoantigen 512; γ-tub – γ-tubulin; R – resting, 2 h, 0 mM glucose; S – stimulated, 25 mM glucose, 2h.

**Suppl. figure S2.**
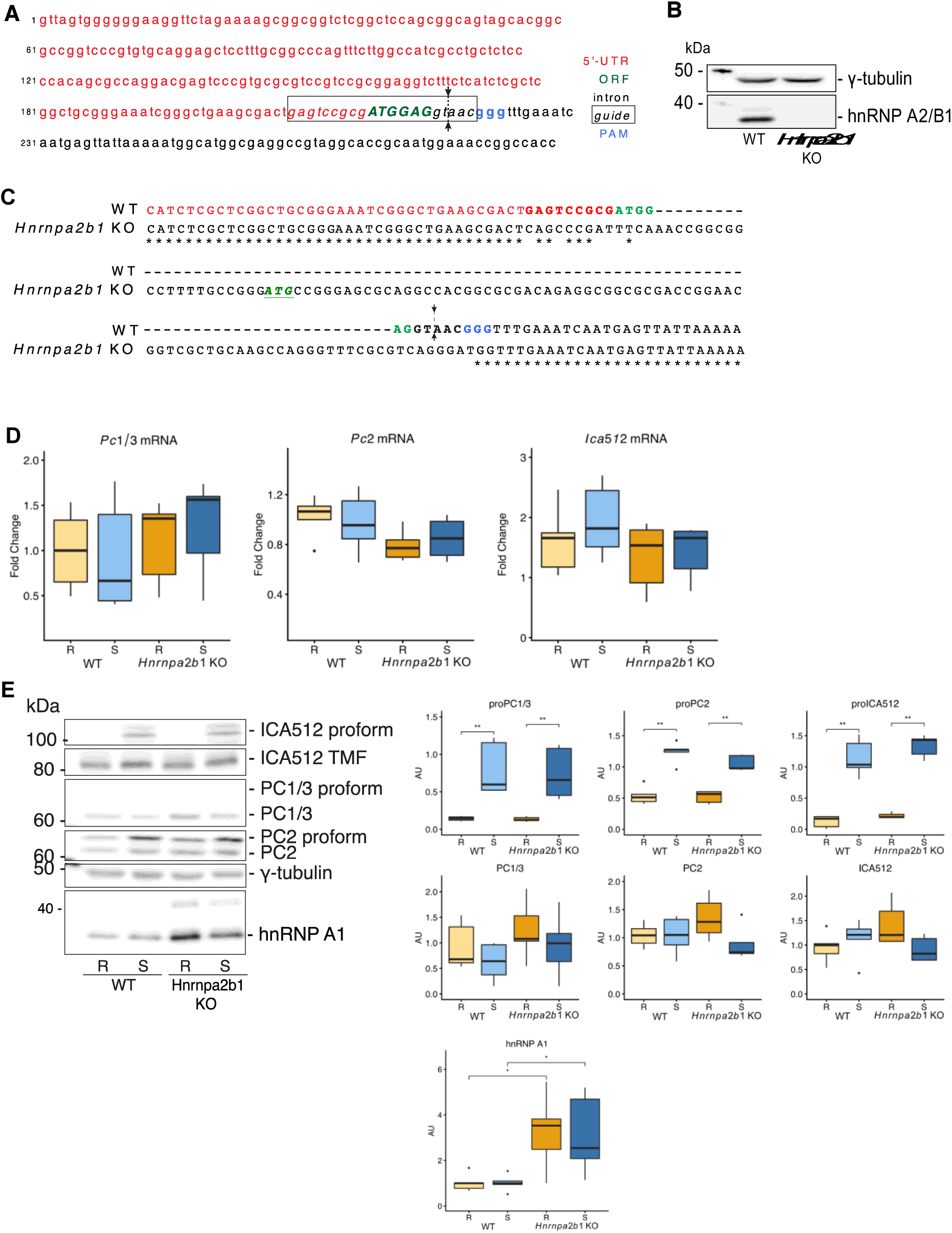
Secretory granule protein expression in *Hnrnpa2b1* KO MIN6 cells. Relates to figure 3. **A:** genomic sequence of the mouse *Hnrnpa2b1* gene. Boxed region defines the guide sequence and arrows show the cut site. **B:** validation of *Hnrnpa2b1* KO MIN6 clone. **C:** sequence comparison between the WT and *Hnrnpa2b1* KO MIN6 cells. Color code as in A. **D:** qPCR of *Pc2*, *Pc1/3* and *Ica512* RNA in resting and stimulated *Hnrnpa2b1* KO MIN6 cells shows no change in mRNA levels compared to WT cells. Normalized to *β-actin* mRNA. **E:** PC2, PC1/3, ICA512 are unchanged in resting and stimulated WT and *Hnrnpa2b1* KO MIN6 cells, while hnRNP A1 levels are increased upon *Hnrnpa2b1* KO. γ-tubulin used as a loading control. Ins – insulin; PC – prohormone convertase; ICA512 – islet cell autoantigen 512; R – resting, 2 h, 0 mM glucose; S – stimulated, 25 mM glucose, 2h. Plots show Tukey-style boxplots with five independent replicates. Mann-Whitney test: * = p<0.05, ** = p<0.01.

**Suppl. Figure S3.**
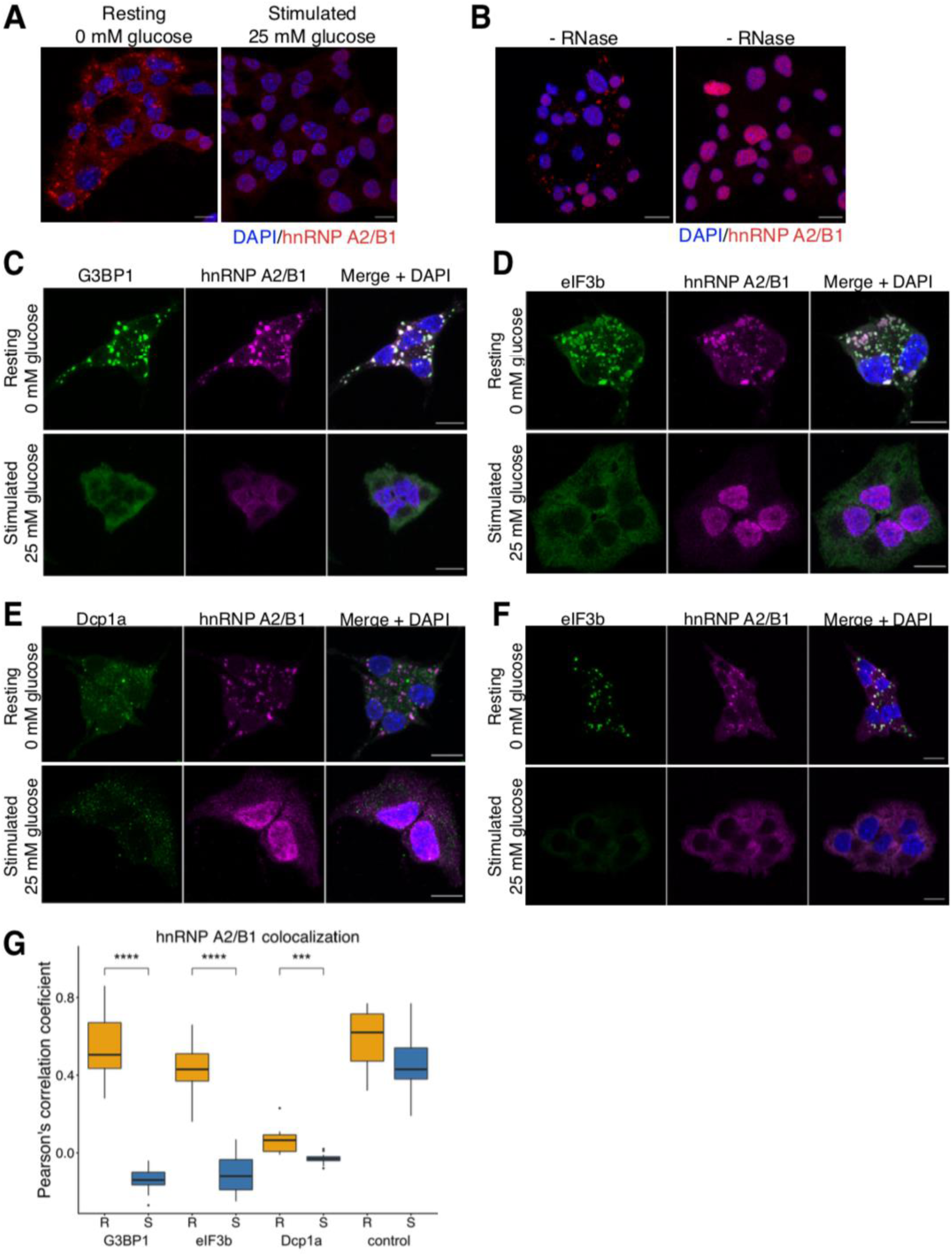
hnRNP A2/B1 localizes to stress granules in resting MIN6 cells. Relates to figure 5 and suppl. figure 5. **A**: hnRNP A2/B1 localizes to RNA granules in cytoplasm of resting MIN6 cells, upon stimulation it is translocated to the nucleus. **B:** resting MIN6 cells were permeabilized with 0.05% saponin, treated with RNase A prior to fixation and stained against hnRNP A2/B1. **C, D, E:** hnRNP A2/B1 colocalizes with stress granule markers G3BP1 (**C**) and eIF3b (**D**) in resting MIN6 cells but not with P body maker Dcp1a (**E**). **F:** resting and stimulated MIN6 cells co-stained with G3BP1 and eIF3b as positive colocalization control. Nuclei were counterstained with DAPI, maximum intensity projections are shown. Scale bar = 10 µm. **G:** Pearson’s correlation coefficient of panels C – F from five independent experiments represented as Tukey-style boxplots; Mann-Whitney test: *** = p<0.005, ** = p<0.01, R – resting, 2 h, 0 mM glucose; S – stimulated, 25 mM glucose, 2h.

**Suppl. Figure S4.**
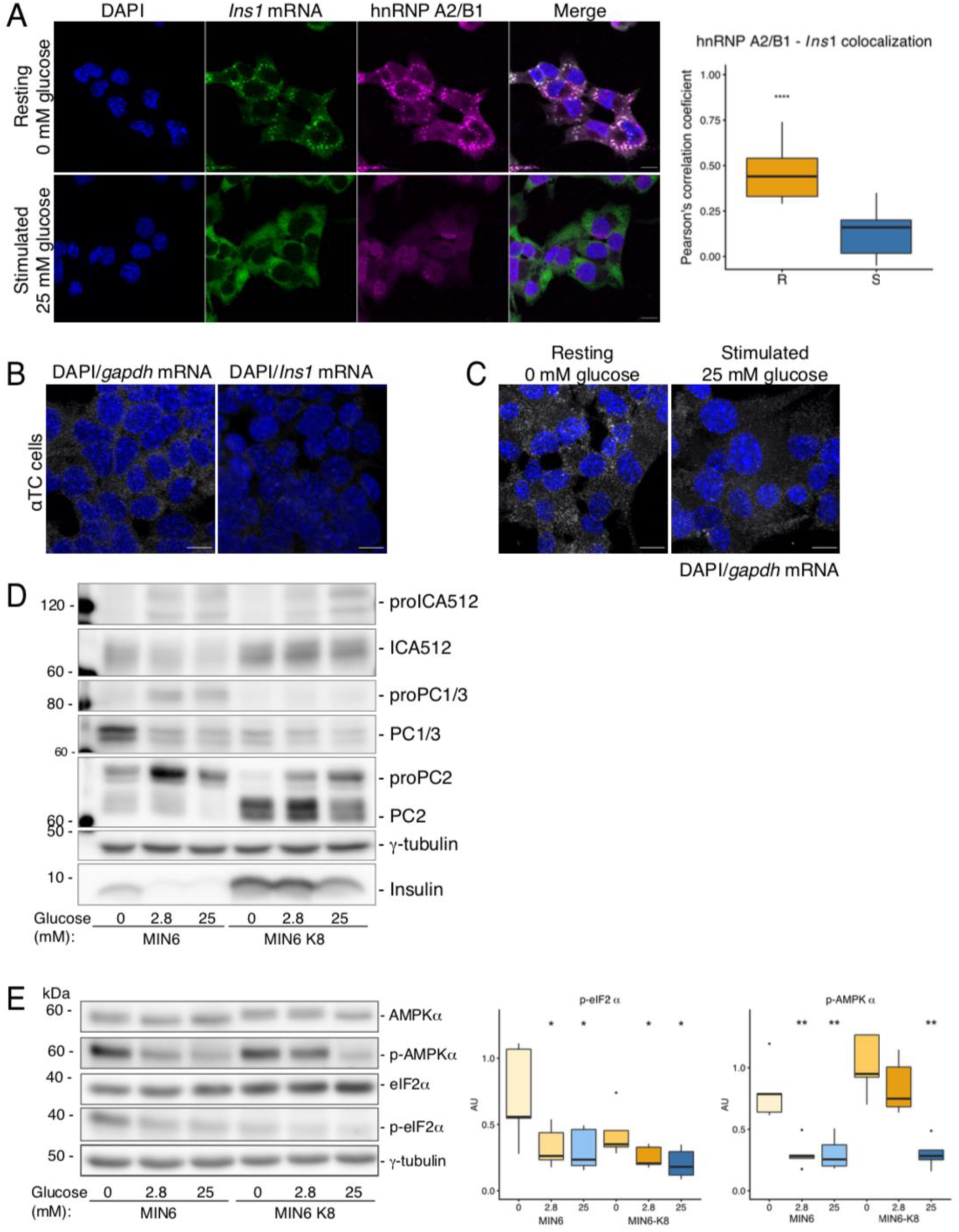
The *Ins1* mRNA colocalizes to hnRNP A2/B1 positive stress granules in MIN6 cells. Relates to figure 5 and suppl. figure 4. **A:** resting MIN6 cells contain hnRNP A2/B1^+^ and *Ins1* mRNA^+^ stress granules. Stimulation disperses the *Ins1* mRNA and hnRNP A2/B1 is translocated to the nucleus. Nuclei were counterstained with DAPI, maximum intensity projections are shown. Scale bar = 10 µm. Pearson’s correlation coefficient of five independent experiments represented as Tukey-style boxplots; t-test: **** = p<0.001; R – resting, 2 h, 0 mM glucose; S – stimulated, 25 mM glucose, 2h. **B:** α-TC cells stained for *Gapdh* (positive control) or *Ins1* mRNA demonstrate probe specificity. Nuclei counterstained with DAPI, scale bar = 10 µm. **C:** Resting and stimulated MIN6 cells stained for *Gapdh* mRNA, nuclei counterstained with DAPI, scale bar = 10 µm. **D:** MIN6 and MIN6-K8 cells differ in upregulation of PC1/2, PC2 and ICA512 proforms in response to different glucose concentrations. **E:** AMPKα and eIF2α phosphorylation in MIN6 and MIN6-K8 cells incubated with 0 mM, 2.8 mM and 25 mM glucose. Ins – insulin; PC2 – prohormone convertase 2; ICA512 – islet cell autoantigen 512; γ-tub – γ-tubulin; R – resting, 2 h, 2.8 mM glucose; S – stimulated, 25 mM glucose, 2h. Plots show Tukey-style boxplots with five independent replicates. Mann-Whitney test compared to MIN6 at 0 mM glucose: * = p<0.05, ** = p<0.01.

**Suppl. Figure S5.**
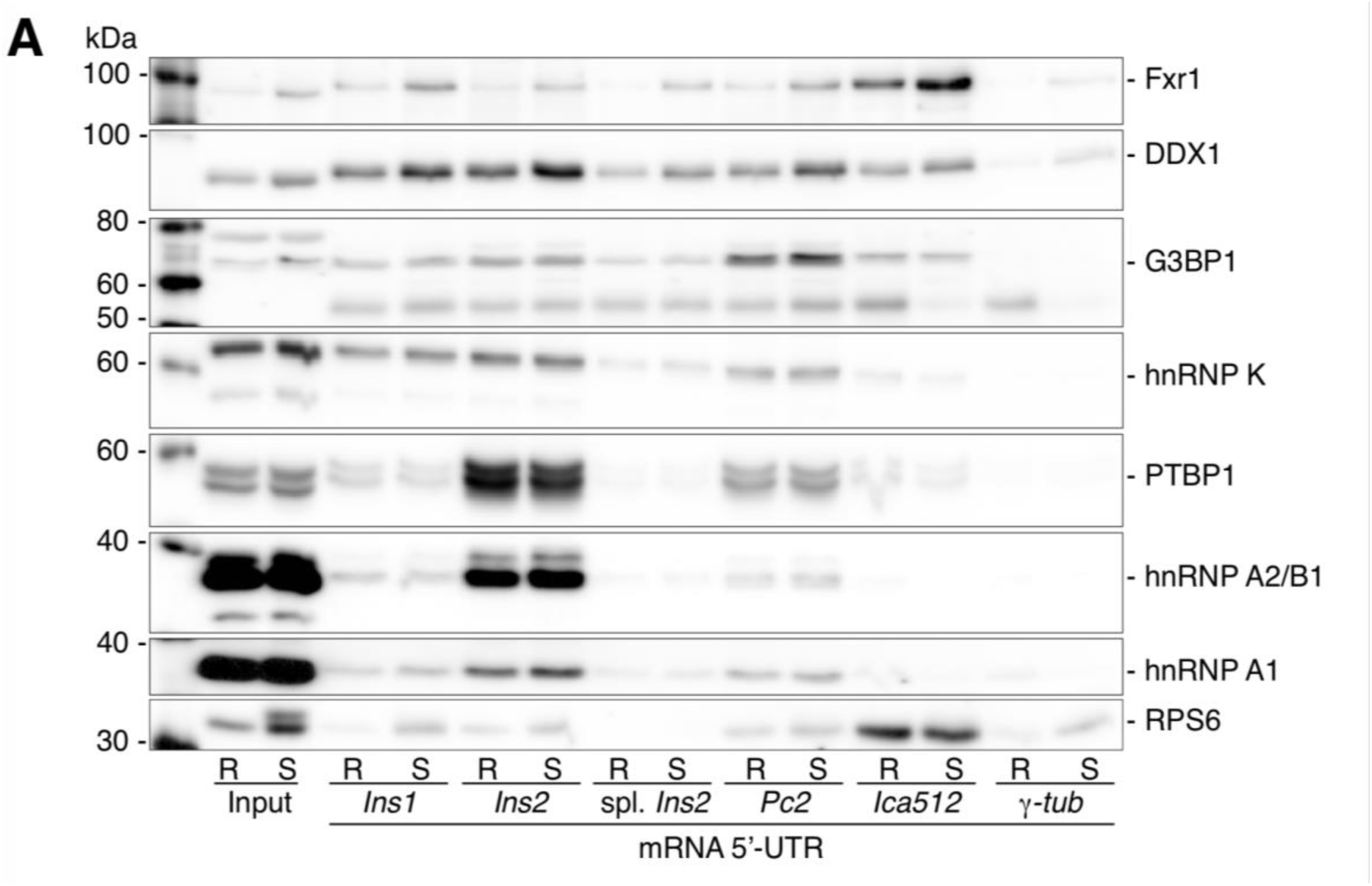
Characterization of the MIN6-K8 clone. Relates to figure 1 and 5, and suppl. figure 1. **A:** RNA pull-downs with the secretory granule protein 5’-UTRs from cytosolic extracts of resting and stimulated MIN6-K8 cells blotted for the RBPs of secretory granule protein RNPs. Ins – insulin; PC2 – prohormone convertase 2; ICA512 – islet cell autoantigen 512; γ-tub – γ-tubulin; R – resting, 2 h, 2.8 mM glucose; S – stimulated, 25 mM glucose, 2h.

**Suppl. Figure S5.**
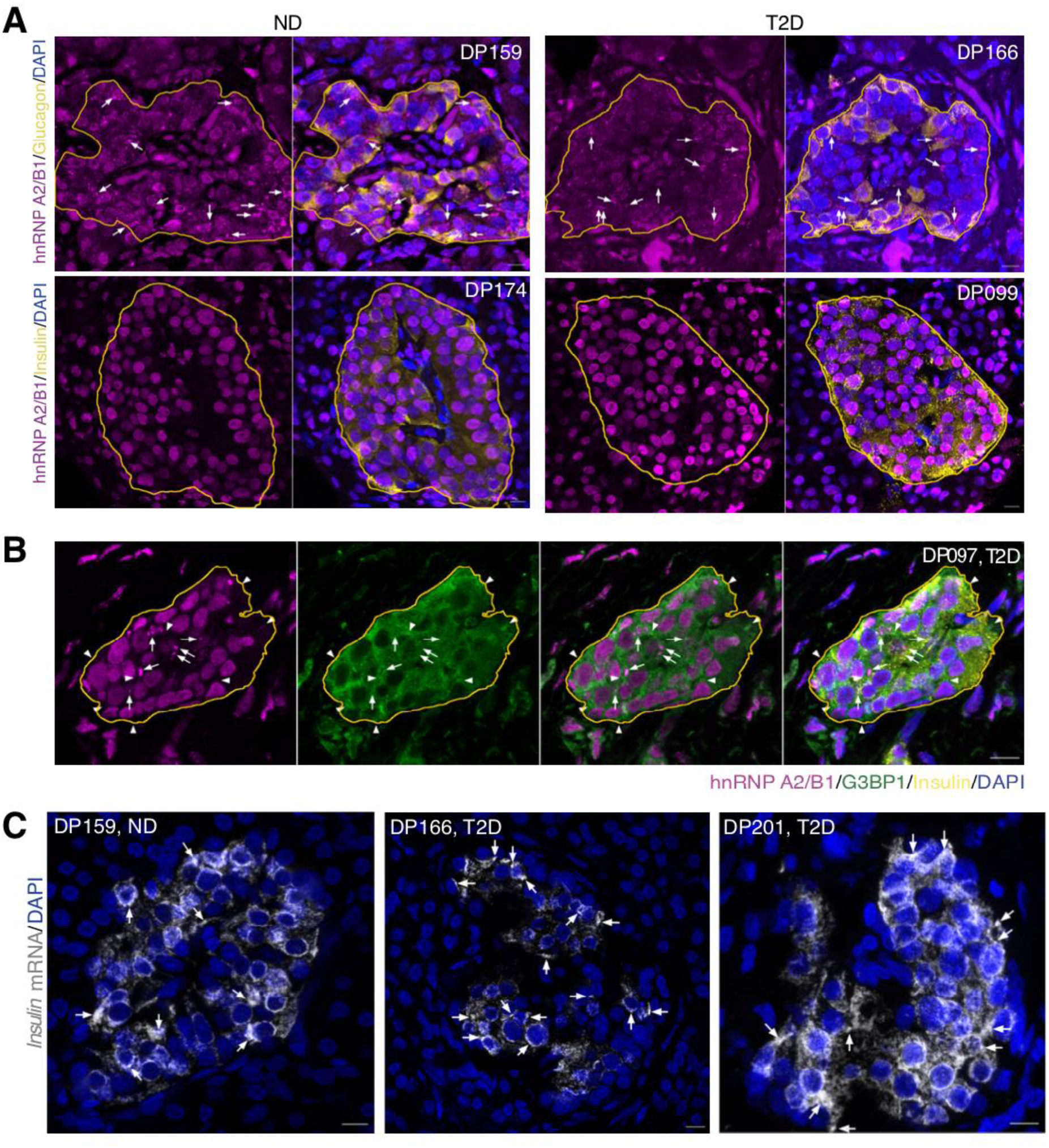
Stress granule markers in human pancreas sections. **A:** FFPE pancreatic sections from partially pancreatomized ND and T2D patients stained with hnRNP A2/B1. Islets are labeled with either insulin or glucagon and circled with yellow lines. Arrows point to cytoplasmic hnRNP A2/B1-positive RNA granules. **B:** FFPE pancreatic sections from partially pancreatomized ND and T2D patients stained with hnRNP A2/B1, G3BP1 and insulin. Islets are circled with yellow lines. Arrows point to cytoplasmic hnRNP A2/B1-positive RNA granules, arrowheads to G3BP1-positive RNA granules. **C:** FFPE pancreatic sections from partially pancreatomized ND and T2D patients stained with the *INS* mRNA. Arrows point to *INS* mRNA-positive RNA granules. Scale bar = 10 µm. ND – non-diabetic; T2D – type 2 diabetic; FFPE – formalin fixed paraffin embedded.

**Suppl. Table 1. Identified proteins identified by MS from *in vitro* pull-downs with the 5’-UTRs of mouse *Ins1, Ins1,* spliced *Ins2, Pc2, Ica512* and *γ-tub* mRNAs and from resting (R) and stimulated (S) MIN6 cells.**

**Suppl. Table 2. Significance analysis of identified proteins.** Significant enrichment of identified proteins was analyzed individually for every target 5’-UTR compared to *γ-tub* 5’-UTR in resting (R) and stimulated (S) conditions using permutation-based t-tests in Perseus, with an FDR of 0.05 and slope s0 of 0.1.

## References

1. Henquin, J.-C. Glucose-induced insulin secretion in isolated human islets: Does it truly reflect β-cell function in vivo? Mol. Metab. 48, 101212 (2021).

2. Ivanova, A. et al. Age-dependent labeling and imaging of insulin secretory granules. Diabetes 62, 3687–96 (2013).

3. Boland, B. B., Rhodes, C. J. & Grimsby, J. S. The dynamic plasticity of insulin production in β-cells. Molecular Metabolism 6, 958–973 (2017).

4. Skelly, R. H., Bollheimer, L. C., Wicksteed, B. L., Corkey, B. E. & Rhodes, C. J. A distinct difference in the metabolic stimulus-response coupling pathways for regulating proinsulin biosynthesis and insulin secretion that lies at the level of a requirement for fatty acyl moieties. Biochem. J. 331 (Pt 2), 553–61 (1998).

5. Mehran, A. E. et al. Hyperinsulinemia drives diet-induced obesity independently of brain insulin production. Cell Metab. 16, 723–737 (2012).

6. Tillmar, L., Carlsson, C. & Welsh, N. Control of insulin mRNA stability in rat pancreatic islets: Regulatory role of a 3′-untranslated region pyrimidine-rich sequence. J. Biol. Chem. 277, 1099–1106 (2002).

7. Schuit, F. C., Kiekens, R. & Pipeleers, D. G. Measuring the balance between insulin synthesis and insulin release. Biochem. Biophys. Res. Commun. 178, 1182–1187 (1991).

8. Guest, P. C., Rhodes, C. J. & Hutton, J. C. Regulation of the biosynthesis of insulin-secretory-granule proteins. Co-ordinate translational control is exerted on some, but not all, granule matrix constituents. Biochem. J. 257, 431–7 (1989).

9. Wicksteed, B. et al. Cooperativity between the Preproinsulin mRNA Untranslated Regions Is Necessary for Glucose-stimulated Translation. J. Biol. Chem. 276, 22553–22558 (2001).

10. Davidson, H. W. & Hutton, J. C. The insulin-secretory-granule carboxypeptidase H. Purification and demonstration of involvement in proinsulin processing. Biochem. J. 245, 575–82 (1987).

11. Molinete, M., Irminger, J.-C., Tooze, S. A. & Halban, P. A. Trafficking/sorting and granule biogenesis in theβ-cell. Semin. Cell Dev. Biol. 11, 243–251 (2000).

12. Lemaire, K., Chimienti, F. & Schuit, F. Zinc transporters and their role in the pancreatic β-cell. J. Diabetes Investig. 3, 202–211 (2012).

13. Itoh, N. & Okamoto, H. Translational control of proinsulin synthesis by glucose [33]. Nature 283, 100–102 (1980).

14. Welsh, M., Nielsen, D. A., MacKrell, A. J. & Steiner, D. F. Control of insulin gene expression in pancreatic beta-cells and in an insulin-producing cell line, RIN-5F cells. II. Regulation of insulin mRNA stability. 260, 13590–13594 (1985).

15. Guest, P. C., Bailyes, E. M., Rutherford, N. G. & Hutton, J. C. Insulin secretory granule biogenesis. Co-ordinate regulation of the biosynthesis of the majority of constituent proteins. Biochem. J. 274 (**Pt 1**, 73–78 (1991).

16. Knoch, K. P. et al. Polypyrimidine tract-binding protein promotes insulin secretory granule biogenesis. Nat. Cell Biol. 6, 207–214 (2004).

17. Gerstberger, S., Hafner, M. & Tuschl, T. A census of human RNA-binding proteins. Nat. Rev. Genet. 15, 829–845 (2014).

18. Singh, G., Pratt, G., Yeo, G. W. & Moore, M. J. The Clothes Make the mRNA: Past and Present Trends in mRNP Fashion. Annu. Rev. Biochem. 84, 325–54 (2015).

19. Kilchert, C., Sträßer, K., Kunetsky, V. & Änkö, M. From parts lists to functional significance— RNA–protein interactions in gene regulation. WIREs RNA 11, (2020).

20. Youn, J. Y. et al. Properties of Stress Granule and P-Body Proteomes. Molecular Cell 76, 286–294 (2019).

21. Advani, V. M. & Ivanov, P. Stress granule subtypes: an emerging link to neurodegeneration. Cellular and Molecular Life Sciences 1–19 (2020). doi:10.1007/s00018-020-03565-0

22. Tauber, D., Tauber, G. & Parker, R. Mechanisms and Regulation of RNA Condensation in RNP Granule Formation. Trends in Biochemical Sciences (2020). doi:10.1016/j.tibs.2020.05.002

23. Hammonds, P., Schofield, P. N., Ashcroft, S. J. H., Sutton, R. & Gray, D. W. Regulation and specificity of glucose-stimulated insulin gene expression in human islets of Langerhans. FEBS Lett. 223, 131–137 (1987).

24. Muralidharan, B., Bakthavachalu, B., Pathak, A. & Seshadri, V. A minimal element in 5′UTR of insulin mRNA mediates its translational regulation by glucose. FEBS Lett. 581, 4103–4108 (2007).

25. Wicksteed, B. et al. A cis-Element in the 5′ Untranslated Region of the Preproinsulin mRNA (ppIGE) Is Required for Glucose Regulation of Proinsulin Translation. Cell Metab. 5, 221–227 (2007).

26. Knight, S. W. & Docherty, K. RNA-protein interactions in the 5’ untranslated region of preproinsulin mRNA. J. Mol. Endocrinol. 8, 225–34 (1992).

27. Magro, M. G. & Solimena, M. Regulation of β-cell function by RNA-binding proteins. Molecular Metabolism 2, 348–355 (2013).

28. Uchizono, Y., Alarcón, C., Wicksteed, B. L., Marsh, B. J. & Rhodes, C. J. The balance between proinsulin biosynthesis and insulin secretion: where can imbalance lead? *Diabetes*, Obes. Metab. 9, 56–66 (2007).

29. Knoch, K. P. et al. cAMP-dependent phosphorylation of PTB1 promotes the expression of insulin secretory granule proteins in β cells. Cell Metab. 3, 123–134 (2006).

30. Knoch, K. P. et al. PTBP1 is required for glucose-stimulated cap-independent translation of insulin granule proteins and Coxsackieviruses in beta cells. Mol. Metab. 3, 518–530 (2014).

31. Süss, C. et al. Rapid changes of mRNA-binding protein levels following glucose and 3-isobutyl-1-methylxanthine stimulation of insulinoma INS-1 cells. Mol. Cell. Proteomics 8, 393–408 (2009).

32. Ehehalt, F. et al. Impaired insulin turnover in islets from type 2 diabetic patients. Islets 2, 30–36 (2010).

33. Fred, R. G., Bang-Berthelsen, C. H., Mandrup-Poulsen, T., Grunnet, L. G. & Welsh, N. High Glucose Suppresses Human Islet Insulin Biosynthesis by Inducing miR-133a Leading to Decreased Polypyrimidine Tract Binding Protein-Expression. PLoS One 5, e10843 (2010).

34. Heni, M. et al. Polymorphism rs11085226 in the Gene Encoding Polypyrimidine Tract-Binding Protein 1 Negatively Affects Glucose-Stimulated Insulin Secretion. PLoS One 7, 1–7 (2012).

35. Lee, E. K. et al. RNA-Binding Protein HuD Controls Insulin Translation. Mol. Cell 45, 826–835 (2012).

36. Kim, C. et al. RNA-binding protein HuD reduces triglyceride production in pancreatic β cells by enhancing the expression of insulin-induced gene 1. Biochim. Biophys. Acta - Gene Regul. Mech. 1859, 675–685 (2016).

37. Fred, R. G., Mehrabi, S., Adams, C. M. & Welsh, N. PTB and TIAR binding to insulin mRNA 3′- and 5′UTRs; implications for insulin biosynthesis and messenger stability. Heliyon 2, 1–26 (2016).

38. Li, Z. et al. RNA-binding protein DDX1 is responsible for fatty acid-mediated repression of insulin translation. Nucleic Acids Res. 46, 12052–12066 (2018).

39. Gräwe, C., Stelloo, S., van Hout, F. A. H. & Vermeulen, M. RNA-Centric Methods: Toward the Interactome of Specific RNA Transcripts. Trends in Biotechnology (2020). doi:10.1016/j.tibtech.2020.11.011

40. Wentworth, B. M. et al. The ratio of mouse insulin I:insulin II does not reflect that of the corresponding preproinsulin mRNAs. Mol. Cell. Endocrinol. 86, 177–186 (1992).

41. Tyanova, S. et al. The Perseus computational platform for comprehensive analysis of (prote)omics data. Nature Methods 13, 731–740 (2016).

42. Liu, Y. & Shi, S. L. The roles of hnRNP A2/B1 in RNA biology and disease. Wiley Interdisciplinary Reviews: RNA 12, (2021).

43. Low, Y. H., Asi, Y., Foti, S. C. & Lashley, T. Heterogeneous Nuclear Ribonucleoproteins: Implications in Neurological Diseases. Molecular Neurobiology 58, 631–646 (2021).

44. Ainger, K. et al. Transport and localization elements in myelin basic protein mRNA. J. Cell Biol. 138, 1077–87 (1997).

45. Gao, Y., Tatavarty, V., Korza, G., Levin, M. K. & Carson, J. H. Multiplexed dendritic targeting of α calcium calmodulin-dependent protein kinase II, neurogranin, and activity-regulated cytoskeleton-associated protein RNAs by the A2 pathway. Mol. Biol. Cell 19, 2311–2327 (2008).

46. Kim, H. J. et al. Mutations in prion-like domains in hnRNPA2B1 and hnRNPA1 cause multisystem proteinopathy and ALS. Nature 495, 467–73 (2013).

47. Martinez, F. J. et al. Protein-RNA Networks Regulated by Normal and ALS-Associated Mutant HNRNPA2B1 in the Nervous System. Neuron 92, 780–795 (2016).

48. Munro, T. P. et al. Mutational analysis of a heterogeneous nuclear ribonucleoprotein A2 response element for RNA trafficking. J. Biol. Chem. 274, 34389–95 (1999).

49. Iwasaki, M. et al. Establishment of new clonal pancreatic β-cell lines (MIN6-K) useful for study of incretin/cyclic adenosine monophosphate signaling. J. Diabetes Investig. 1, 137–142 (2010).

50. Kedersha, N. et al. Evidence that ternary complex (eIF2-GTP-tRNAiMet)-Deficient preinitiation complexes are core constituents of mammalian stress granules. Mol. Biol. Cell 13, 195–210 (2002).

51. Szkudelski, T. & Szkudelska, K. The relevance of AMP-activated protein kinase in insulin-secreting β cells: a potential target for improving β cell function? Journal of Physiology and Biochemistry 75, 423–432 (2019).

52. Sonenberg, N. & Hinnebusch, A. G. Regulation of Translation Initiation in Eukaryotes: Mechanisms and Biological Targets. Cell 136, 731–745 (2009).

53. Panas, M. D., Ivanov, P. & Anderson, P. Mechanistic insights into mammalian stress granule dynamics. Journal of Cell Biology 215, 313–323 (2016).

54. Ashcroft, S. J., Bunce, J., Lowry, M., Hansen, S. E. & Hedeskov, C. J. The effect of sugars on (pro)insulin biosynthesis. Biochem. J. 174, 517–26 (1978).

55. Solimena, M. et al. Systems biology of the IMIDIA biobank from organ donors and pancreatectomised patients defines a novel transcriptomic signature of islets from individuals with type 2 diabetes. Diabetologia 61, 641–657 (2018).

56. Barovic, M. et al. Metabolically phenotyped pancreatectomized patients as living donors for the study of islets in health and diabetes. Molecular Metabolism 27, S1–S6 (2019).

57. Farack, L. et al. Transcriptional Heterogeneity of Beta Cells in the Intact Pancreas. Dev. Cell 48, 115–125.e4 (2019).

58. Kulkarni, S. D. et al. Glucose-stimulated translation regulation of insulin by the 5’ UTR-binding proteins. J. Biol. Chem. 286, 14146–56 (2011).

59. Kanai, Y., Dohmae, N. & Hirokawa, N. Kinesin transports RNA: Isolation and characterization of an RNA-transporting granule. Neuron 43, 513–525 (2004).

60. Elvira, G. et al. Characterization of an RNA granule from developing brain. Mol. Cell. Proteomics 5, 635–651 (2006).

61. Jain, S. et al. ATPase-Modulated Stress Granules Contain a Diverse Proteome and Substructure. Cell 164, 487–498 (2016).

62. Smith, R. Moving molecules: mRNA trafficking in mammalian oligodendrocytes and neurons. Neuroscientist 10, 495–500 (2004).

63. Wigger, L. et al. Multi-omics profiling of living human pancreatic islet 1 donors reveals heterogeneous beta cell trajectories 2 toward type 2 diabetes 3 4. bioRxiv 2020.12.05.412338 (2020). doi:10.1101/2020.12.05.412338

64. Patel, A. et al. A Liquid-to-Solid Phase Transition of the ALS Protein FUS Accelerated by Disease Mutation. Cell 162, 1066–1077 (2015).

65. Douglas, J., Gardner, L., Salapa, H. & Levin, M. Antibodies to the RNA Binding Protein Heterogeneous Nuclear Ribonucleoprotein A1 Colocalize to Stress Granules Resulting in Altered RNA and Protein Levels in a Model of Neurodegeneration in Multiple Sclerosis. J. Clin. Cell. Immunol. 07, 402 (2016).

66. Mandrioli, J., Mediani, L., Alberti, S. & Carra, S. ALS and FTD: Where RNA metabolism meets protein quality control. Seminars in Cell and Developmental Biology (2019). doi:10.1016/j.semcdb.2019.06.003

67. Xue, Y. C. et al. Dysregulation of RNA-Binding Proteins in Amyotrophic Lateral Sclerosis. Frontiers in Molecular Neuroscience 13, (2020).

68. Clarke, J. P. W. E. et al. Multiple sclerosis-associated hnrnpa1 mutations alter hnrnpa1 dynamics and influence stress granule formation. Int. J. Mol. Sci. 22, 1–22 (2021).

69. Tsuboi, T., Da, G., Xavier, S., Leclerc, I. & Rutter, G. A. 5-AMP-activated Protein Kinase Controls Insulin-containing Secretory Vesicle Dynamics* □ S. (2003). doi:10.1074/jbc.M307800200

70. Leclerc, I. et al. Metformin, but not leptin, regulates AMP-activated protein kinase in pancreatic islets: Impact on glucose-stimulated insulin secretion. Am. J. Physiol. - Endocrinol. Metab. 286, (2004).

71. Pepin, É. et al. Pancreatic β-cell dysfunction in diet-induced obese mice: Roles of amp-kinase, protein kinase Cɛ, mitochondrial and cholesterol metabolism, and alterations in gene expression. PLoS One 11, 153017 (2016).

72. Brun, T. et al. AMPK profiling in rodent and human pancreatic beta-cells under nutrient-rich metabolic stress. Int. J. Mol. Sci. 21, (2020).

73. Del Guerra, S. et al. Functional and molecular defects of pancreatic islets in human type 2 diabetes. Diabetes 54, 727–735 (2005).

74. Vasiljević, J., Torkko, J. M., Knoch, K. P. & Solimena, M. The making of insulin in health and disease. Diabetologia 63, 1981–1989 (2020).

## Additional references

75. Phelps, E. A. et al. Advances in pancreatic islet monolayer culture on glass surfaces enable super-resolution microscopy and insights into beta cell ciliogenesis and proliferation. Sci. Rep. 7, (2017).

76. Xie, F., Xiao, P., Chen, D., Xu, L. & Zhang, B. miRDeepFinder: A miRNA analysis tool for deep sequencing of plant small RNAs. Plant Mol. Biol. 80, 75–84 (2012).

77. Cox, J. & Mann, M. MaxQuant enables high peptide identification rates, individualized p.p.b.- range mass accuracies and proteome-wide protein quantification. Nat. Biotechnol. 26, 1367–1372 (2008).

78. Cox, J. et al. Accurate proteome-wide label-free quantification by delayed normalization and maximal peptide ratio extraction, termed MaxLFQ. Mol. Cell. Proteomics 13, 2513–2526 (2014).

